# A frontal motor circuit for economic decisions and actions

**DOI:** 10.64898/2025.12.11.693624

**Authors:** Oliver M. Gauld, Chaofei Bao, Jingjie Li, Gauthier Boeshertz, Timothy P. H. Sit, Joseph Warren, Joseph O. Tutt, Yang Pan, Nikolaos Zervogiannis, Claudia Clopath, Jeffrey C. Erlich, Chunyu A. Duan

**Author notes:** These authors contributed equally (Oliver M. Gauld, Chaofei Bao).

## Abstract

Flexible behaviour requires transforming abstract cognitive representations, such as value preferences, into concrete motor actions. During economic decision-making, individuals evaluate options to guide choices and then transform these choices into specific actions to obtain rewards. Understanding how neural circuits convert these abstract economic decisions into spatial actions remains challenging because decision formation and motor planning are typically intertwined. Here we introduce a mouse task that temporally dissociates value-guided decisions from spatial action planning, and show that a frontal motor network implements the transformation across decision stages through dynamic circuit reconfiguration. Using cortex-wide imaging and optogenetic perturbations, we identified a frontal motor circuit that was causally required for both abstract and motor stages of choice. During the abstract decision stage, neurons in this circuit encoded option values and economic choices independently of sensorimotor contingencies, and unilateral silencing impaired decisions without spatial bias. In contrast, during spatial planning, value and spatial signals were non-linearly integrated to guide action selection, and unilateral silencing produced an ipsilateral bias. A dynamical model captured this transformation, predicting a mode switch from cooperative interhemispheric maintenance of economic choice to competitive, lateralized control of actions, which we validated with simultaneous bilateral recordings. These findings demonstrate how frontal motor circuits reconfigure their interactions to bridge abstract cognition and concrete actions, providing a circuit-level mechanism for flexible, value-guided behaviour.

## Introduction and Results

In daily life, actions are guided by individual decision preferences which shape subjective utility, a measure of internal value and satisfaction. This process is broadly referred to as economic decision-making. Choices between different options, such as food sources or financial investments, often require integrating information across multiple dimensions, including reward size^1^, probability^2,3^, time delay^4,5^, and effort^6,7^. Understanding how economic choices are formed based on integrated option values is critical for explaining adaptive behaviours as well as maladaptive choice patterns seen in disorders such as gambling addiction, obsessive compulsive disorder, and impaired financial decision-making^8,9^. Multiple theoretical frameworks have been proposed to formally describe how the brain transforms option values into actions. According to the good-based model, an abstract decision is made by comparing different option values independent of sensorimotor contingencies^10^, and this abstract choice then flexibly guides action selection. In contrast, the action-based model posits that values are assigned directly to competing actions, such that decision formation and motor planning unfold together^11,12^.

Neural signatures of abstract value representation have been found in the primate orbitofrontal cortex (OFC)^13^ among other regions, in support of the good-based model. However, value-modulated action selection signals have also been observed across multiple cortical and subcortical areas^14–18^, consistent with the action-based model. Hybrid models propose that option and action selection proceed in parallel^19^, reflecting the reciprocal interactions between value and motor systems. Reconciling these frameworks remains challenging for two reasons. First, in most decision-making tasks, decision formation and action planning occur simultaneously, making it difficult to isolate the underlying computations. Second, the types of neural signals observed depend strongly on the brain regions examined. Addressing the fundamental question of how abstract cognitive representations are transformed into concrete actions therefore requires task designs that temporally separate decision and action processes (Fig. 1a) and systematic cross-region comparisons. A rare example of such dissociation was achieved in a primate study, where neural signatures of value-to-action transformation were found in lateral prefrontal cortices, whereas only abstract value signals were found in OFC^20^. Nevertheless, the underlying circuit mechanisms and causal contributions to such transformation remain unknown. Recent advances in rodent models of value-based decision-making provide an opportunity to conduct causal, circuit-level dissection of economic choices^21–28^.

**Fig. 1:**
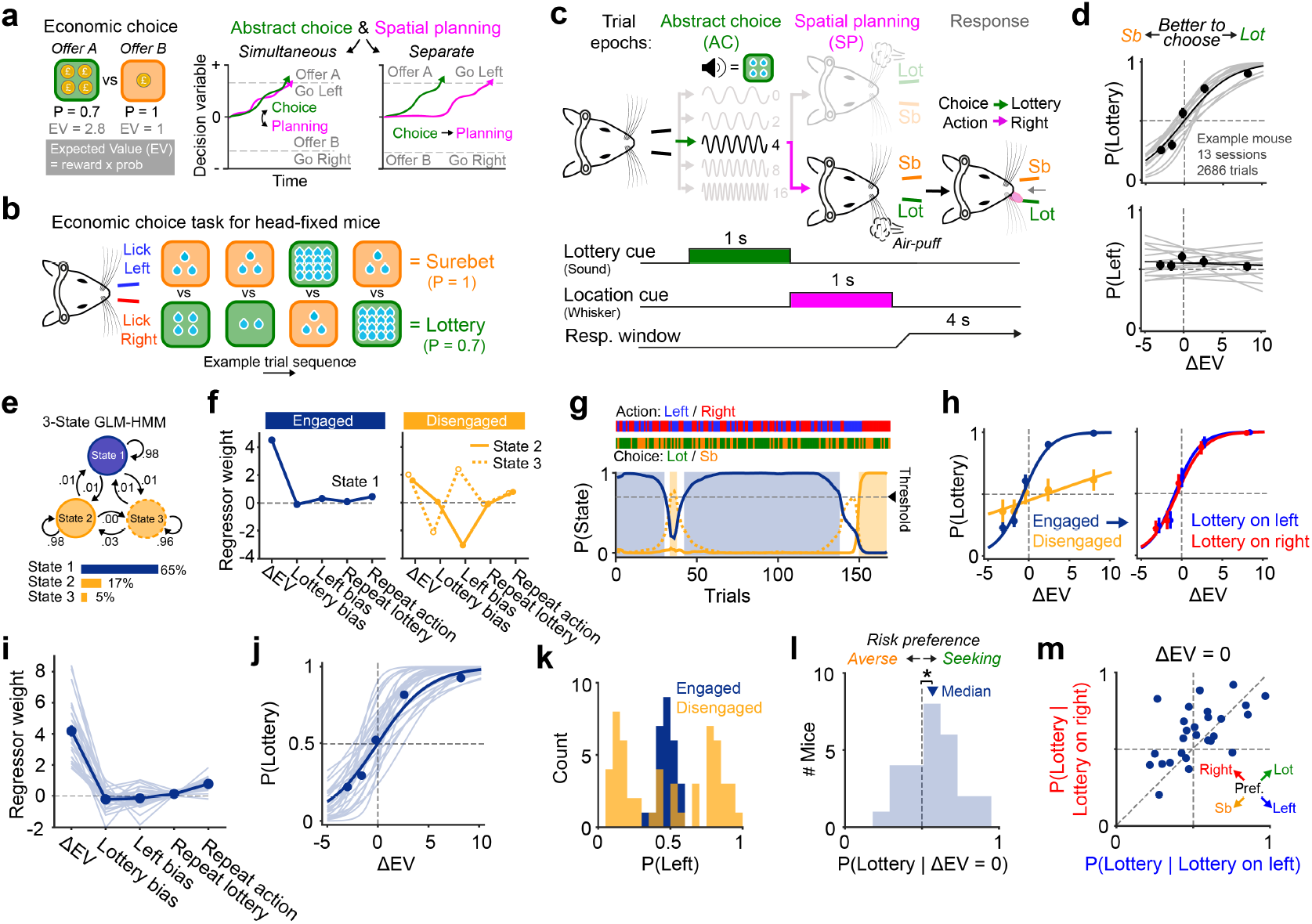
Economic decision-making task design and performance. **a**, Typical task design where abstract economic choice and spatial action planning occur simultaneously; versus our task design with temporally separate decision stages. **b**, Example trial sequence. Lottery offer value and spatial location were randomised across trials. Mice indicated choice with directional licking. **c**, Stimuli and trial timing. **d**, Behavioural performance of an example mouse. Top, lottery choice probability as a function of Δ*EV* = − *EV*_*lottery*_ *EV*_*surebet*_. Bottom, leftward choice probability as a function of ΔEV. Grey, individual sessions; black, concatenated data across sessions. **e**, Example 3-state GLM-HMM for one mouse. The numbers and arrows indicate transition probabilities. The bars indicate the percent of trials assigned to each state with P(State) ≥ 0.7. **f**, Inferred GLM weights for the 3 states shown in **e. g**, Posterior state probabilities for an example session. The horizontal dashed lines denotes the P(State) threshold (0.7). Colour code as in **f. h**, Left, lottery choice preference shown separately for engaged and disengaged trials for an example mouse. Right, engaged trials split by lottery offer location. **i**, Inferred GLM weights during the engaged state across mice. Data points indicate mean ± s.e.m.; individual lines represent individual mice (n = 27). **j**, Behavioural performance on engaged trials across mice. **k**, Spatial bias during engaged and disengaged states across mice. **l**, Distribution of risk preference during engaged trials across mice, two-tailed *t*-test against 0.5, n = 27 mice, *P* = 0.0439. **m**, Risk preference as a function of lottery location (left vs. right). Individual points represent mice.

Towards this goal, we developed a novel task in head-fixed mice with distinct temporal epochs that separate abstract economic choice from spatial action planning within each trial. Combining mesoscale imaging and bilateral optogenetic perturbations across the dorsal cortex, we identified a frontal motor circuit, specifically the anterolateral motor cortex (ALM), that is causally required for both economic decisions and subsequent actions. High-density electrophysiological recordings showed that ALM neurons encode option values and abstract economic choice independent of sensorimotor contingencies during the initial decision epoch, before representing spatial planning and execution signals in later epochs^29–32^. To further investigate how abstract decision signals in ALM are transformed into spatial motor plans, we employed unilateral perturbations, dynamical modelling and simultaneous recording in both hemispheres. We found a mode switch in interhemispheric interactions across decision formation and action planning stages, from cooperative to competitive, providing a circuit-level mechanism for value-to-action transformation during economic decision-making.

### A novel paradigm to study economic decisions and actions

We trained mice on a novel task designed to dissociate value-guided decision-making from preparatory motor processes associated with behavioural report (Fig. 1a-c; Fig. S1; Methods). On each trial, head-fixed mice chose between two offers: a guaranteed “surebet” reward (3 µL; P(reward) = 1) or a probabilistic “lottery” (0 − 16 µL; P(reward) = 0.7). The expected value (EV), defined as the product of reward magnitude and probability (Fig. 1a), varied across trials for lottery offers and was cued by an auditory stimulus (Fig. 1b,c). To maximize reward, choices should be guided by the EV difference (ΔEV) between the surebet and current lottery offer. Following the value cue, a left or right whisker stimulation indicated the spatial locations of lottery and surebet offers on that trial (Fig. 1c). For example, right whisker stimulation signalled that the lottery was on the right, prompting a right lick for lottery and left lick for surebet. Auditory and whisker cues were randomised across trials so that neither lottery magnitude nor location could be predicted. Stimulus contingencies were stable within each mouse but counterbalanced across mice (Fig. S2a). After an initial learning period, mice’s lottery choice preferences were guided by ΔEV with min-imal spatial biases (Fig. 1d, Fig. S1b), and performance and response time (RT) patterns were consistent across contingency groups (Fig. S2b,e).

To control for fluctuations of task engagement over time, we fit a hidden Markov model (HMM) where behavioural performance was characterized by probabilistic transitions between latent states (Fig. 1e), each associated with a distinct weighting of task-relevant generalized linear model (GLM) regressors (Fig. 1f)^33,34^. Mice’s performance was well captured by a 3-state GLM-HMM (Fig. S3a,b), allowing us to distinguish trials where lottery choices were predominantly guided by ΔEV (‘engaged’) from those driven by internal or history-dependent biases (‘disengaged’; Fig. 1f,g, Fig. S3c,d). Psychometric curves of an example expert animal showed that choices depended strongly on offer values but not on offer locations in the engaged state (Fig. 1h). These findings are consistent across individual mice (Fig. 1i-k) and generalized across cohorts (Fig. S2c,d). Task engagement peaked mid-session (Fig. S3e), revealing an inverted-U relationship between optimal performance and time-varying factors such as satiation, impulsivity and attention. Risk preference in the engaged state, defined as lottery choice probability when offer EVs were equal, varied across individuals (Fig. 1l), but was reliable between left and right lottery locations (Pearson’s *r* = 0.58, *P* = 1.8 × 10^−3^; Fig. 1m). Taken together, these behavioural results show that mice can integrate temporally separate value and spatial information to flexibly guide lottery versus surebet choices during active task engagement, establishing a foundation for investigating the neural mechanisms underlying the value-to-action transformation. From here on, we refer to the non-spatial, value-guided decisions between lottery versus surebet offers as “abstract economic choices” (see Discussion); and preparation for upcoming left versus right licking responses as “spatial action planning”.

### Distributed and state-dependent task representation on the dorsal cortex

To characterize how dorsal cortical activity represents key components of the task across distinct stages of economic decision-making, we performed widefield calcium imaging in transgenic mice expressing GCaMP6s in excitatory neurons (n = 6 expert mice, 8.82 ± 1.72 sessions per mouse, 246.46 ± 65.12 trials per session, mean ± std; Fig. 2a); while monitoring facial, body movements and pupillometry using videography (Fig. 2b, Fig. S4). We used a linear encoding model^35^ to relate cortex-wide dynamics to task events and movement variables (Fig. 2c). Task event regressors included lottery value cue, whisker stimulation side, abstract (lottery vs surebet) choice, and spatial (left vs right) choice. Data were concatenated across sessions for each mouse, and subject-level models were fit before averaging results across mice. The model reliably predicted single-trial activity with high overall cross-validated performance (Fig. 2d).

**Fig. 2:**
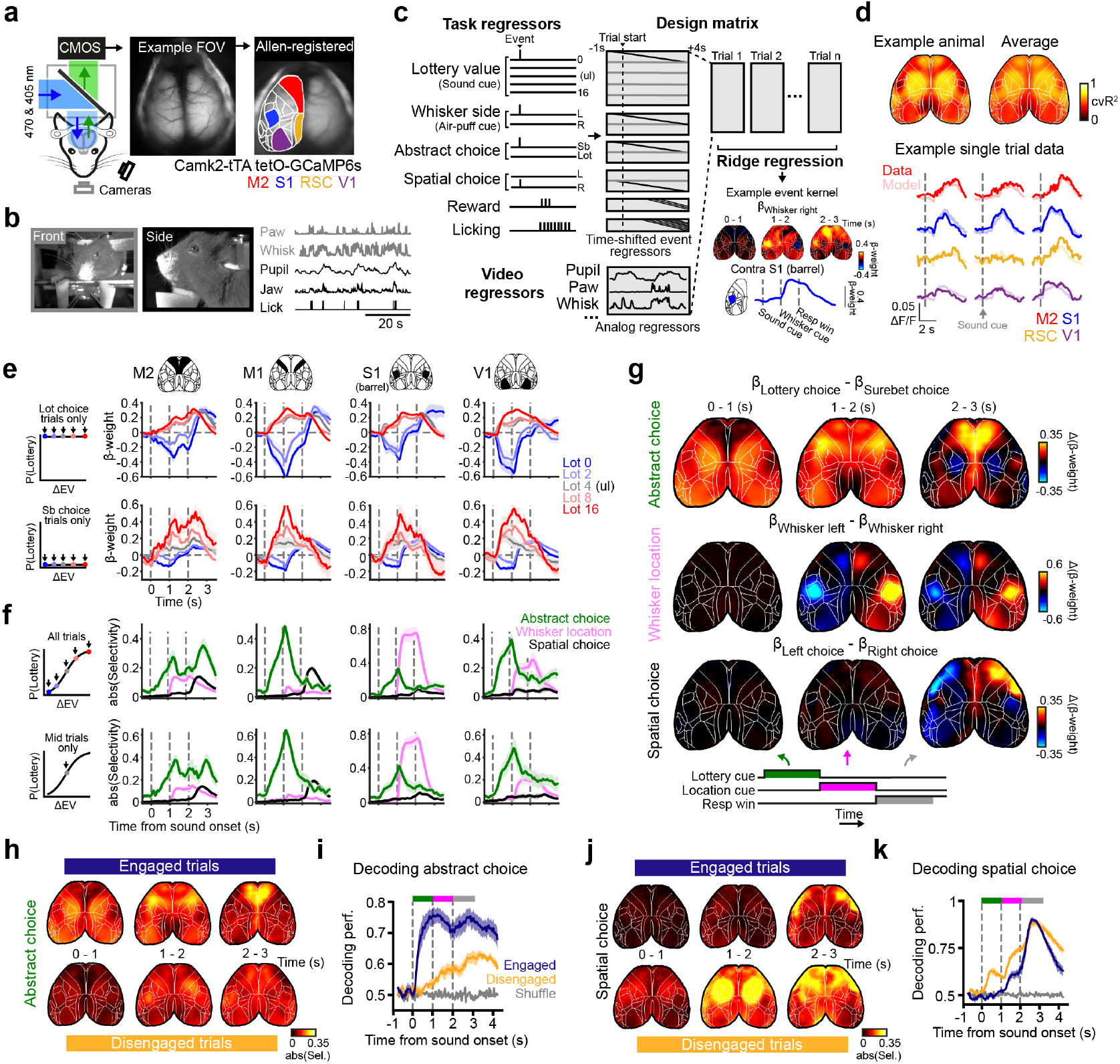
Spatiotemporal representations of task information across the dorsal cortex. **a**, Left, widefield calcium imaging (WF) set-up. Middle, dorsal cortex field-of-view (FOV) imaged at 470 nm. Right, FOV aligned to the Allen Common Coordinate Framework (CCF) with example areas highlighted. **b**, Example behavioural videography images and extracted movement signals. **c**, Ridge regression model design. An example pixel-wise kernel map for the right air-puff regressor is shown for a single animal averaged across trial epochs. The accompanying trace shows the mean kernel response from left barrel cortex. **d**, Model performance. Top, pixel-wise maps showing cross-validated model explained variance across the whole trial for an example mouse (left) and averaged across mice (right; n = 6 mice). Bottom, single-trial fluorescence (dark), and model-predicted activity (light) across different cortical areas for three example trials. **e**, Lottery value encoding. Kernel regressor traces for the five lottery trial types averaged across different bilateral cortical areas. Top, model trained on lottery choice trials; bottom, model trained on surebet choice trials. **f**, Selectivity traces for abstract choice (green), whisker location (pink) and spatial choice (black) averaged across bilateral cortical regions. Selectivity was computed as the absolute difference between the respective paired event kernel regressors. Top, model trained on all trials; bottom, model trained only on middle lottery value trials. **g**, Pixel-wise signed selectivity maps averaged across trial epochs. Top, abstract choice; middle, whisker location; bottom, spatial choice. **h**, Pixel-wise selectivity maps for abstract choice for engaged (top) and disengaged (bottom) trials. **i**, Decoding of abstract choice using fluorescence activity across all CCF-defined cortical areas across engaged and disengaged trials. **j**, Similar to **h**, but for spatial choice. **k**, Similar to **i**, but for spatial choice. Data in **e**-**k** show mean ± s.e.m across 6 mice, 52 sessions, 7804 engaged trials, 2401 disengaged trials. Data in **e**-**g** show results from engaged trials.

Spatiotemporal profiles of the event kernels revealed distributed cortical representation of task variables. Following the lottery cue, activity across multiple cortical regions correlated positively with lottery value, even after controlling for economic choice (Fig. 2e). This value-related activity was consistent in mice with opposite mappings between auditory tone frequencies and lottery values (Fig. S5b), confirming that this signal reflects value rather than sensory features. In addition to value coding, we observed distributed representation of the upcoming abstract choice, with higher activity for lottery compared to surebet choices (Fig. 2f green, g top). To verify that abstract choice coding existed independent of offer value coding, we focused on trials of a single lottery type at the midpoint of the psychometric curve, where ΔEV was close to 0 and mice had similar probability of lottery and surebet choices (Fig. 2f bottom). We found that even when mice received the exact same sensory stimulus, cortical activity was still predictive of abstract choices. Because this abstract choice signal appeared in the first epoch, when only offer values but not offer locations were available, it indicates that mice indeed computed abstract economic decisions before spatial action planning. In regions such as the secondary motor cortex (M2), this selectivity persisted into later action planning, execution, and reward epochs. Abstract choice signals were also modulated by task engagement (Fig. 2h). Choice decoding accuracy was higher and emerged earlier during engaged than disengaged states (Fig. 2i; Fig. S5c,g), revealing state modulation of decision strategies and the underlying neural representations. Activity was higher for lottery than surebet choices across all mice, regardless of whether the whisker stimulus cued the lottery or surebet location (Fig. S5d), suggesting that these signals likely encode value-guided decisions rather than a sensorimotor rule of “licking toward or away from the upcoming whisker stimulation”.

In contrast to the distributed value and economic choice representations, spatial signals were more localized. Whisker stimulation activated contralateral barrel and whisker motor cortices (Fig. 2f pink, g middle row), and was similar across the engaged and disengaged states (Fig. S5e,h). Spatial choice activity emerged during the second epoch, when offer value and location information were combined to guide actions (Fig. 2f black, k), with strong lateralised action signals in motor and somatosensory cortex during the execution of directional licking (Fig. 2g bottom). Notably, during disengaged states, spatial choice could be decoded even in the first epoch (Fig. 2j,k; Fig. S5i), reflecting disengaged choices driven by internal biases rather than by integrating task-relevant stimuli. Decoding sensory, decision and action variables from the entire dorsal cortex yielded higher accuracies than decoding from single regions (Fig. S5f), suggesting that some task information is encoded in patterns of multi-region dynamics. Overall, mesoscale imaging revealed widespread, state-dependent representation of economic choices and actions across the dorsal cortex, with regions such as M2 containing all task-relevant information in corresponding decision stages.

### Causal roles of frontal motor cortex for economic decisions and actions

To test whether the observed cortical representations are required for task performance, we conducted a causal survey of the dorsal cortex using bilateral optogenetic inhibition in transgenic mice expressing Channelrhodopsin-2 (ChR2) in inhibitory interneurons (VGAT-ChR2-EYFP). In each session, we randomly targeted 17 cortical locations by scanning a blue laser over the intact skull in a subset of trials (Fig. 3a)^30,36^, either during the first or second task epoch (1 s). Electrophysiological recordings in awake mice confirmed that laser stimulation effectively silenced cortical activity within a ~ 1.5 to 2 mm radius in a power-dependent manner (Fig. S6a-d)^37,38^, with similar results across cortical areas (Fig. S6c) and minimal rebound after laser offset (Fig. S6e). Deeper cortical and subcortical regions were unlikely to be inhibited due to confinement of light spread and ChR2 expression (Fig. S6d)^30^.

**Fig. 3:**
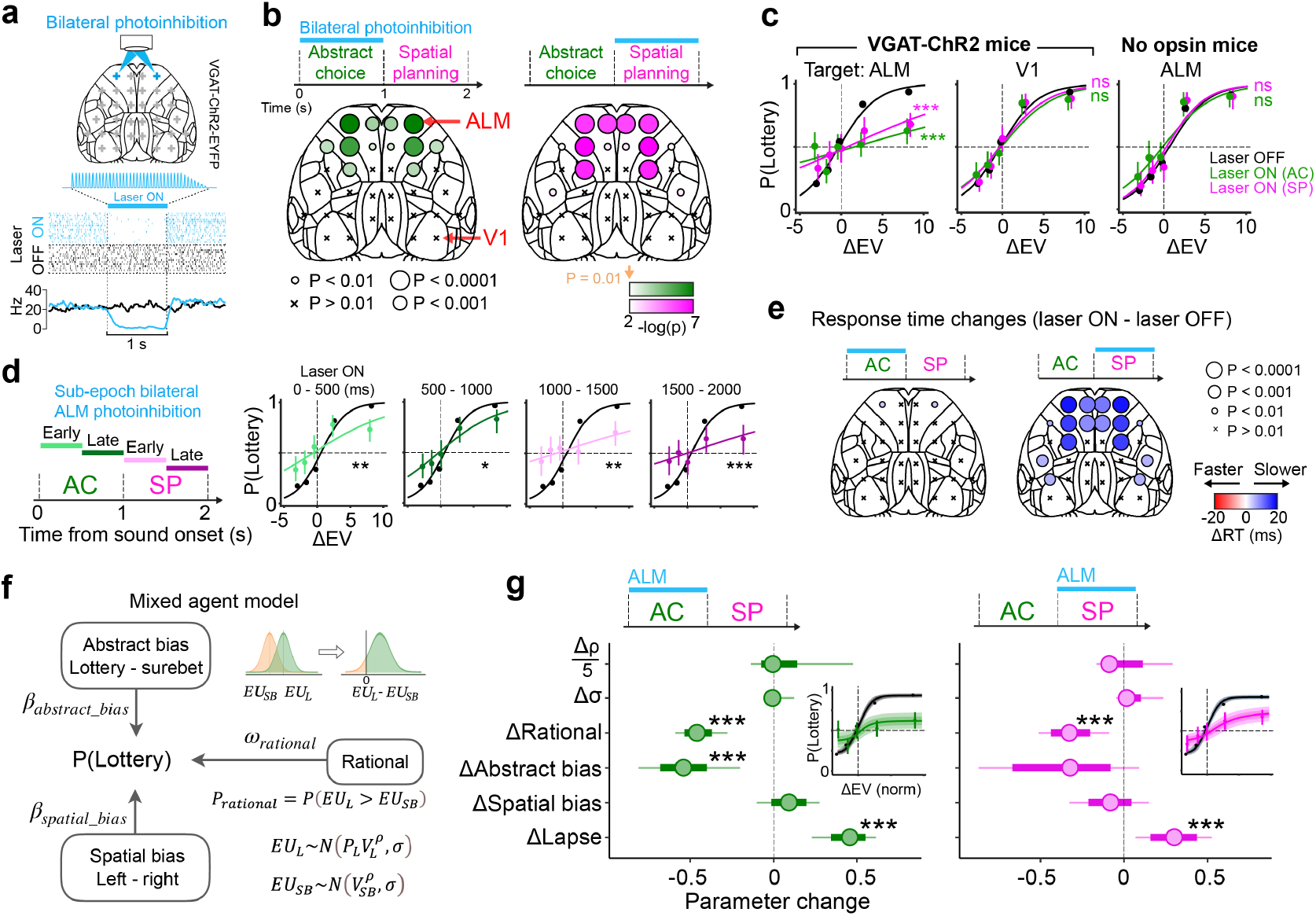
Cortical areas involved in task performance revealed by laser-scanning photoinhibition. **a**, Top, laser-scanning photoinhibition mapping of dorsal cortex in VGAT-ChR2-EYFP mice. Bottom, spike raster and peri-stimulus time histogram (psth) from a putative pyramidal neuron in ALM on 4 mW laser ON (blue) and OFF (black) trials. **b**, Behavioural effects of photoinhibition. Epoch and region-specific effects of photoinhibition on lottery choices were quantified using generalised linear mixed-effects models (GLMMs). *P* values were corrected for multiple comparisons (17 regions). **c**, Psychometric curves during epoch-specific photoinhibition. Left, ALM inhibition in VGAT-ChR2 mice (n = 8); middle, V1 inhibition in VGAT-ChR2 mice (n = 8); right, ALM laser delivery in control mice (n = 6). Black, laser OFF; green and magenta, photoinhibition during the first and second epochs, respectively. Data show mean and 95% binomial confidence intervals (CI) across trials. **d**, Psychometric curves for sub-epoch ALM photoinhibition. Data show mean and 95% binomial CI. **e**, Response time (RT) changes during bilateral photoinhibition. **f**, Mixed-agent model schematic. *EU*, expected utility; *P*, lottery payout probability; *V*, lottery and surebet reward magnitude; *ρ* controls utility curvature; *σ* controls decision noise. **g**, Left, mixed-agent model fits to bilateral ALM photoinhibition during the abstract choice epoch, parameter changes (Δ) indicate perturbation effects compared to control performance. Right, same as Left but for spatial planning epoch photoinhibition. Inset shows model psychometric fits to the behaviour data. Statistical notes: In **b-d**, statistical significance was assessed using cross-validated (CV) GLMM model comparison. The median *P* value across CV repetitions is shown. In **e**, paired comparisons between RTs on laser ON versus laser OFF (control) trials used Wilcoxon ranksum tests. *P* values were corrected for multiple comparisons (17 regions). In **g**, two-tailed significance was tested using posterior confidence intervals. Significance is indicated as \**P* < 0.05, \*\**P* < 0.01, \*\*\**P* < 0.001; ns, non-significant *P* ≥ 0.05.

In contrast to the widespread cortical representations of offer value and economic choice revealed by mesoscale imaging (Fig. 2), only a small network of frontal motor regions was causally required during the abstract decision epoch (Fig. 3b left; Fig. S7a-c). We compared lottery choice preference on interleaved laser ON and laser OFF trials during active task engagement (Methods) and quantified the effect of each targeted region using cross-validated model comparison between nested generalized linear mixed-effects models (GLMMs), corrected for multiple comparisons. The largest and most consistent effects occurred after silencing the anterolateral motor cortex (ALM, Fig. 3c left) in M2, which remained significant with weaker, more spatially confined inhibition (Fig. S7d,e), and was consistent across cohorts with varied stimulus contingencies (Fig. S7f). Control mice without ChR2 expression showed no behavioural effects (Fig. 3c, right; Fig. S7b), confirming that performance impairments were not due to light delivery alone. Consistent with previous findings^30^, ALM and neighbouring frontal motor regions were also required for planning directional licking responses in the second epoch (Fig. 3b right, c), even when such planning was embedded in this complex task context. To further examine the temporal specificity of ALM’s causal roles, we delivered 500 ms inactivations targeting early or late phases of the abstract decision and spatial planning epochs, and found that all sub-epoch manipulations significantly impaired performance (Fig. 3d). In addition, photoinhibition during the spatial planning epoch increased response times (RT), whereas inhibition during the abstract choice epoch caused minimal slowing (Fig. 3e; Fig. S7g), suggesting dissociable contributions to preparatory processes.

Although the GLMM analyses are effective for identifying which brain areas are causally required for task performance, they do not provide mechanistic insights into the specific contributions these brain areas make to the decision process. To better understand the perturbation effects, we used a mixed-agent model^21^ that distinguishes whether perturbations impaired value-based decision-making or shifted stimulus-independent biases (Fig. 3f). The model describes baseline behavioural strategies by mixing different types of agents, whose relative influence on choice is determined by the mixing weights, *ω*. The first agent is a ‘rational’ agent that maximizes expected utility, a measure of subjective value that considers offer values as well as individual animals’ risk preference (controlled by *ρ*) and decision noise (*σ*). The other agents are stimulus-independent agents that either habitually choose the lottery or the surebet (abstract bias) or habitually choose the left or right (spatial bias). Residual random choices are captured by a lapse parameter. Model validation using synthetic data confirmed accurate recovery of generative parameters (Fig. S8a).

To characterize how optogenetic silencing shifted the relative influence of each agent/strategy compared to control performance, we estimated the joint posterior over the parameters for laser ON and OFF trials across animals. Bilateral ALM inhibition during either the abstract choice or spatial planning epochs significantly reduced rational decision-making (Fig. 3g; ΔRational ≪ 0) with a corresponding increase in lapse (ΔLapse ≫ 0). Abstract choice epoch inhibition additionally shifted the abstract bias towards the habitual surebet agent (ΔAbstract bias ≪ 0). These effects were absent following V1 inactivation and in control mice lacking ChR2 expression (Fig. S8d,e). Together with the value coding revealed by widefield imaging, the model-based quantifications of optogenetic silencing identify the frontal motor network, especially ALM, as a key circuit required for rational, value-guided choice.

### Mixed economic value, choice, and action coding in ALM neurons

To examine dynamic coding of task variables at cellular resolution, we performed high-density electrophysiological recordings in ALM of trained mice (n = 4 mice; 2446 cells; 17 sessions; Fig. S9a). We found heterogeneous tuning for spatial action planning (Fig. 4a,b Example neuron 1), option value (Fig. 4a,b Example neuron 2), abstract choice, and mixed selectivity (Fig. 4a,b Example neuron 3) across decision stages. During the first epoch, nearly half of ALM neurons (46.5%) showed significant modulation by lottery value, non-spatial abstract (lottery vs surebet) choice, or both (Fig. 4c). Among value-selective neurons, most fired more strongly for high-than low-value options (503 versus 190, *chi*^2^ = 141.0, *P* = 3 × 10^−32^); and most choice-selective neurons had higher firing rates for lottery than surebet choices (533 versus 222, *chi*^2^ = 128.2, *P* = 3 × 10^−29^). Although option value and abstract choice selectivity were correlated across the population (*r* = 0.48, *P* = 7.44 × 10^−144^), we confirmed that value representation was present when accounting for abstract choices and vice versa (Methods). Note that our task design encourages, but does not enforce, abstract decision formation before planning a spatial action. Mice could, in principle, maintain offer values in working memory without forming an explicit abstract decision^39^, and directly perform value-guided action selection when offer locations are revealed. Our recordings from ALM support that abstract economic decisions were made in the first epoch, independent of the spatial configurations of offers or upcoming lick directions (see Discussion).

**Fig. 4:**
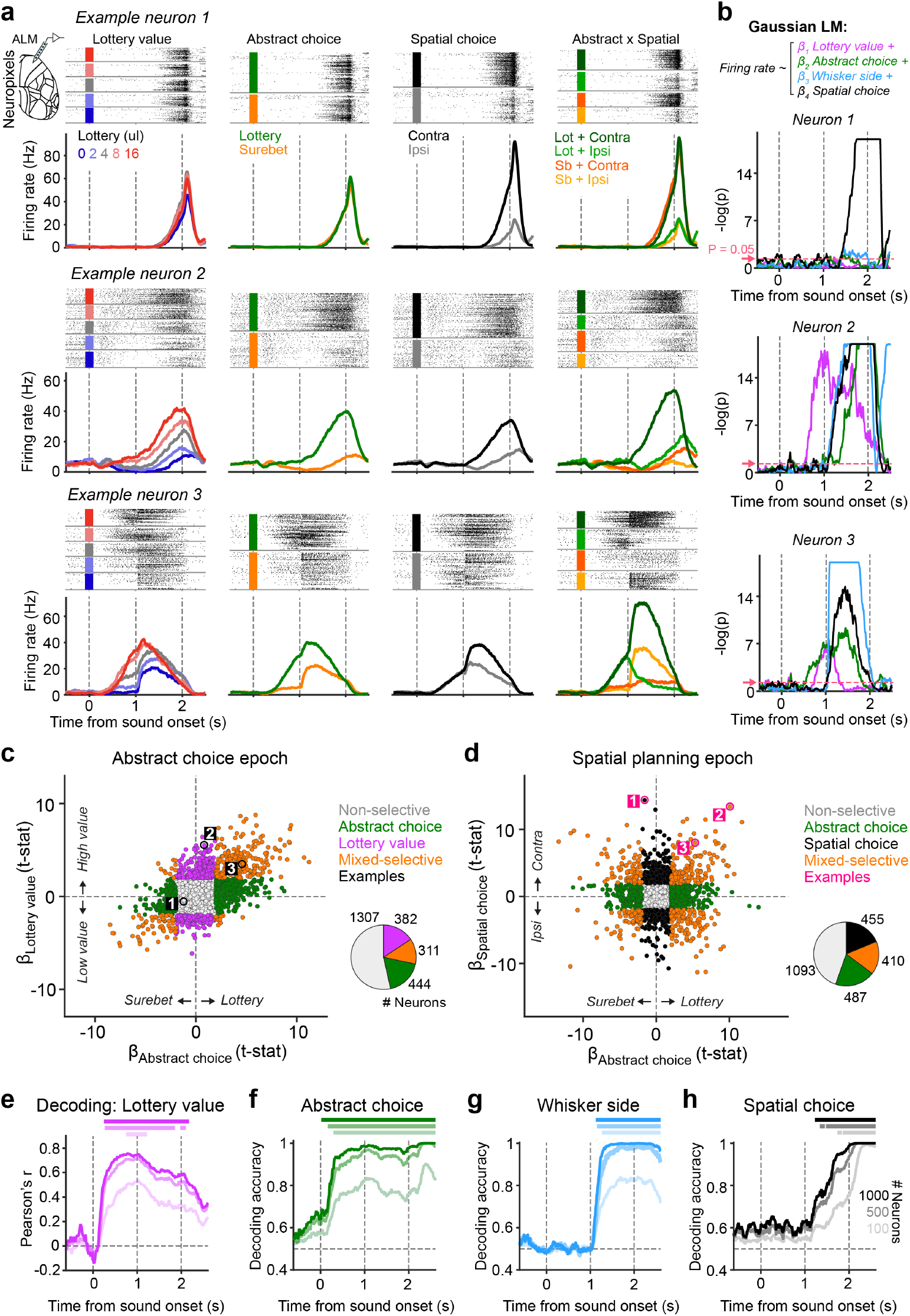
Neural signatures of value-to-action transformation in ALM. **a**, Spike rasters and PSTHs from three example neurons recorded in ALM using Neuropixels probes. Neural activity is aligned to the sound cue onset and sorted across trial conditions: lottery offer values (left), abstract lottery versus surebet choices (middle left), spatial choices contra- or ipsilateral to the recording side (middle right), and combinations of abstract and spatial choice conditions (right). **b**, Linear model of single-neuron activity. Significance of each regressor is shown across time for the three example neurons in **a**. The arrows and horizontal dashed lines indicate *P* = 0.05. **c**, Coefficients of lottery value versus abstract choice regressors during the first epoch (0 − 1 s). Each point represents an individual neuron (example neurons marked in black); colours indicate significantly selective neurons (*P* < 0.05). Inset pie chart summarizes the number and proportion of selective neurons. **d**, Coefficients of spatial choice vs abstract choice regressors during the second epoch (1 − 2 s), similar to **c. e**, Population decoding performance of lottery value over time, for pseudopopulation sizes of 1000 (darkest shade), 500, and 100 (lightest shade) neurons. Decoding accuracy is quantified as the correlation between cross-validated linear model predictions and true lottery values. Spikes are aligned to the sound cue onset and counted over windows of 200 ms with 10 ms shifts between neighbouring windows. Performance is plotted over the right edge of the window (causal). Horizontal bars on top mark the time points with significant decoding accuracy (median *P* < 0.01 across 200 pseudopopulations, Fig. S10). **f**, Population decoding performance for linear classification of abstract choices over time. **g**, Classification performance to linearly separate whisker stimulus sides. **h**, Classification performance to linearly separate spatial choices.

Once offer locations were revealed, ALM dynamics reflected the transformation from value-guided decisions to spatial actions. As illustrated in an example neuron (Fig. 4a Example neuron 3), offer location and abstract choice information were non-linearly multiplexed, mapping abstract choices to directional licking responses. As a result, this neuron represented all task variables during the second epoch (Fig. 4b bottom). Across the population, 35.4% of neurons encoded upcoming spatial actions, and 36.7% encoded abstract choices. These two forms of selectivity were independent (*r* = 0.005, *P* = 0.81; Fig. 4d, Fig. S9b,c), consistent with the orthogonal structure of our task design. A small subset of ALM neurons (16.8%) exhibited conjunctive coding for both abstract and spatial decisions, representing an integrative signal that may underlie the task computation. Comparisons across layers showed significantly larger fractions of task-selective neurons in deep layers of ALM during the first and second epochs (Fig. S9d,e). Among task-selective neurons, ALM deep layers also contained a higher proportion of conjunctive neurons with mixed selectivity for abstract and spatial decisions (38.1%) compared to superficial layers (19.6%; *chi*^2^ = 28.2, *P* = 1.1 × 10^−7^; Fig. S9d,e right). The single-neuron analyses reveal that ALM represents all the task variables required to make the abstract economic choice in the first epoch, and then integrate that choice with whisker input to generate a spatial action plan.

To better estimate the strength and latency of these signals on the population level, we performed cross-validated linear decoding analyses. ALM showed robust information of option value and abstract choice, which emerged early in the first epoch and persisted through spatial planning, movement execution and reward feedback periods (Fig. 4e,f, Fig. S10). Following lateralized whisker stimulation, signals related to offer location and upcoming directional licking responses appeared sequentially in the second epoch (Fig. 4g,h), closely mirroring the computational steps in our task. Increasing pseudopopulation size improved decoding accuracy and reduced latency, indicating distributed coding within ALM. These cellular-resolution recordings confirmed what we found using mesoscale widefield imaging (Fig. 2), and further reveal how single-neuron mixed selectivity enables orthogonal representations of abstract and spatial choices on the population level to support flexible, context-dependent decision-making.

### Unilateral inhibition confirms non-spatial computation for abstract choice

Consistent with previous studies^29–32^, we found spatial planning signals in ALM in the second epoch to guide upcoming directional licking responses. In contrast, ALM’s contribution during abstract decision formation appears non-spatial. To test the lateralized nature of ALM’s causal role across decision stages, we conducted unilateral optogenetic inhibition during either the first or second epoch. We first examined the general impact of inhibition on lottery choice preference, and found a significant reduction of rational decision-making (ΔRational ≪ 0) and a shift towards choosing the surebet (ΔAbstract bias ≪ 0; Fig. 5a,e,f). These effects paralleled those observed with bilateral inhibition (Fig. 3c) but were approximately half in magnitude, suggesting that abstract decisions are maintained through distributed processing across hemispheres. When we separated trials by choice direction relative to the inhibited hemisphere (ipsi- or contralateral), we found a strong ipsilateral response bias (ΔSpatial bias ≫ 0) following second-epoch inhibition but not after first-epoch inhibition (Fig. 5b,e,f). Response times were also significantly slowed after secondepoch but not first-epoch inhibition (Fig. 5c), with stronger effects on contralateral spatial choices (Fig. 5d). Thus, perturbing one side of ALM during the first epoch disrupted abstract economic choices without biasing upcoming directional licking responses; whereas second-epoch inactivation impaired economic choices and biased/slowed spatial responses. No behavioural changes were observed in control mice without ChR2 expression, confirming that these effects were not due to lateralized light delivery (Fig. S8f).

**Fig. 5:**
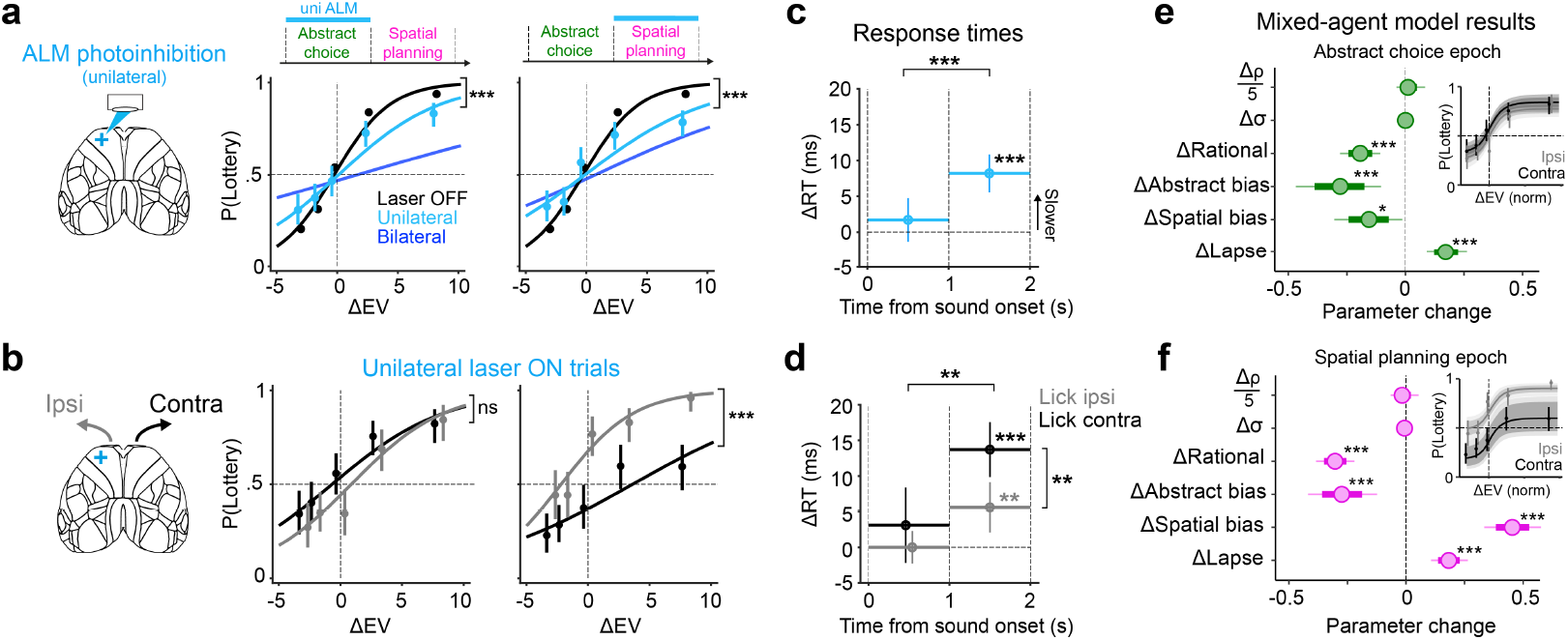
Distinct behavioural effects of unilateral ALM photoinhibition across decision stages. **a**, Unilateral photoinhibition effects on lottery choices for the first (left) or second (right) epoch. Data and psychometric curves are shown for laser OFF (black) and laser ON (light blue) trials. Bilateral inactivation curves are shown for comparison (dark blue). Data points show mean ± 95% binomial CI (n = 8 VGAT-ChR2-EYFP mice). **b**, Epoch-specific optogenetic effects on spatial choice. Unilateral laser ON trials are split by whether the choice was contralateral (black) or ipsilateral (grey) to the inactivated hemisphere. **c**, Average change in response times (RT) following unilateral ALM inactivations. Data points and vertical bars show the median ± s.e.m across trials. The horizontal bars indicate timing and duration of photoinhibition. **d**, Similar to **c**, but showing RT changes split by contralateral (black) or ipsilateral (gray) choices. **e**, Mixed-agent model characterisation of unilateral ALM inactivation during the abstract choice epoch. **f**, Similar to **e**, but for the spatial planning epoch. In **e** and **f**, spatial bias is calculated as the ipsilateral choice bias minus the contralateral choice bias. Only the second epoch inactivation resulted in a strong ipsilateral bias. Statistical notes: In **a**,**b**, significance was assessed using cross-validated (CV) GLMM model comparison. The median *P* value across CV repetitions is shown. In **c**,**d**, paired comparisons across conditions, or comparisons to control trials, used Wilcoxon ranksum tests. In **e**,**f**, two-tailed tests using posterior confidence intervals. Significance is indicated as \**P* < 0.05, \*\**P* < 0.01, \*\*\**P* < 0.001.

### Dynamical model and bilateral ALM recordings test interhemispheric interactions during economic decisions and actions

Why does unilateral ALM inhibition produce a spatial bias in the second but not the first epoch (Fig. 5)? We hypothesized that abstract economic choices are maintained cooperatively by both ALM hemispheres, whereas spatial actions are selected through competitive interactions: the transformation from value-guided decision to lateralized action reflects a dynamic reconfiguration of interhemispheric coordination. To test this idea, we constructed a dynamical model comprising 3 recurrently connected neural populations on each hemisphere: a lottery pool receiving value input positively correlated with ΔEV, a surebet pool receiving value input negatively correlated with ΔEV and mutually inhibitory with the lottery pool, and an action pool that drives movement to the contralateral side (Fig. 6a). During the second epoch, both abstract value-guided pools receive lateralized whisker inputs and project to the contralateral action pool such that lottery choices correspond to licking toward the whisker side. Across hemispheres, abstract choice neurons have cooperative interactions through excitatory connections, whereas spatial action neurons have competitive interactions through inhibitory connections. Abstract decisions thus arise via mutual inhibition between lottery and surebet neurons in both hemispheres; while spatial actions are determined by mutual inhibition between hemispheres.

**Fig. 6:**
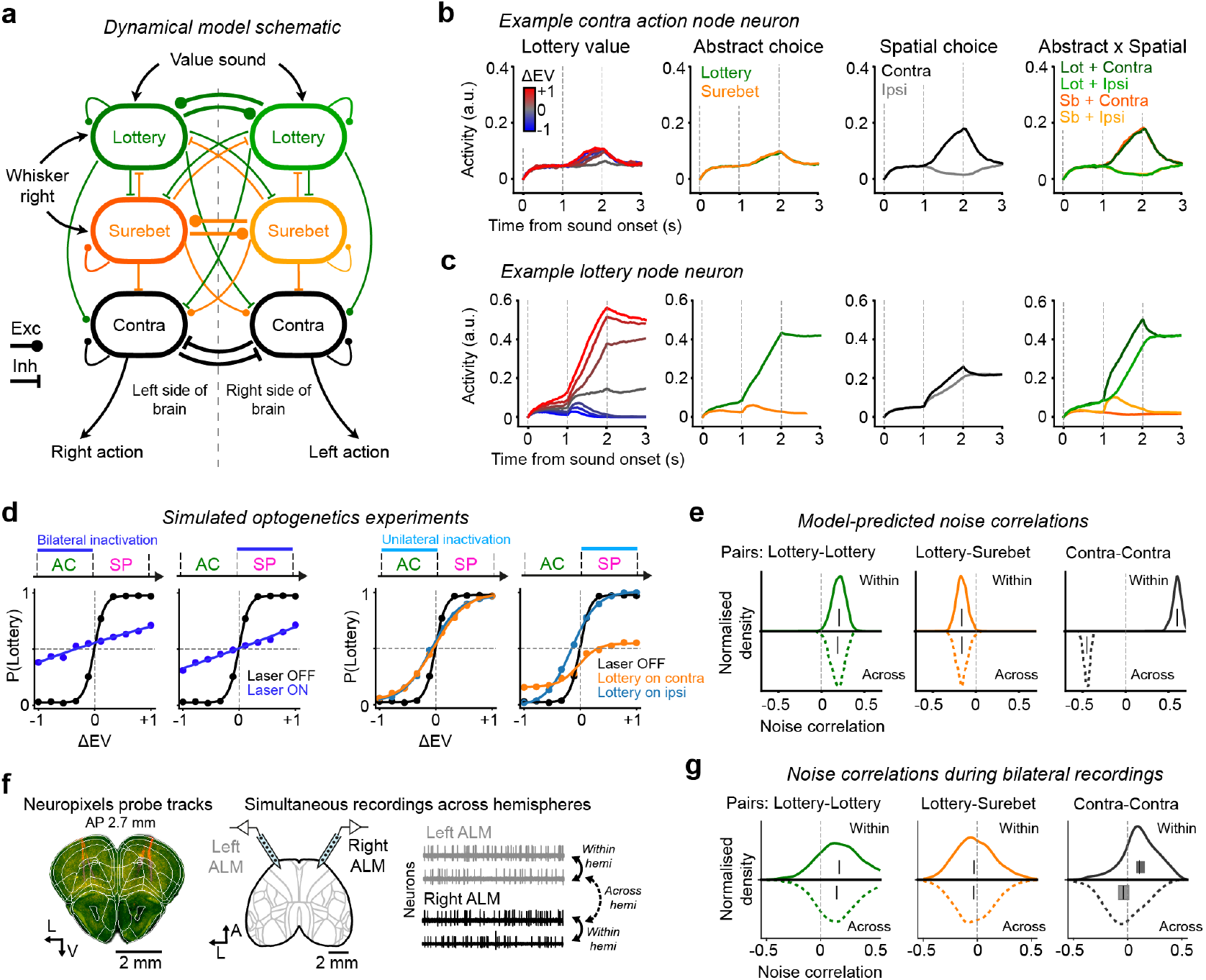
Dynamical ALM model and bilateral recordings support changing interhemispheric interactions across decision stages. **a**, Schematic of circuit model, showing 6 pools of recurrently connected nodes, each pool representing a population of ALM neurons. Abstract value pools have excitatory connections between hemispheres whereas motor pools have inhibitory connections between hemispheres. To avoid overcrowding, not all inputs are shown. See Methods for complete description of the model. **b**, Activity of one example Contra-preferring neuron on the left hemisphere for different trial conditions (averaged over 20 trials per condition), showing contralateral action selectivity in the second epoch. **c**, Activity of one example Lottery-preferring neuron on the left hemisphere, showing value and abstract choice selectivity for both task epochs. **d**, Model simulations qualitatively reproduce ALM bilateral (left) and unilateral (right) inactivation effects during the first and second epoch. Unilateral inactivations during the spatial-planning epoch resulted in a spatial bias. **e**, Noise correlation predictions from the dynamical model, for pairs of neurons that both prefer lottery choices (left), neuron pairs with opposite value tunings (middle), and neuron pairs that drive contralateral choices (right), within each hemisphere (top half, 870 pairs) and across hemispheres (bottom half, 900 pairs). Activity from the first epoch is used for estimating noise correlations between value neurons and activity from the second epoch is used for estimating noise correlations between motor neurons. Kernel density estimation is plotted for each distribution. **f**, Schematic and example histology for simultaneous bilateral ALM recordings using Neuropixels probes. Schematic traces from 4 simultaneously recorded neurons, permitting within and across hemisphere comparison of noise correlations. **g**, Similar to **e**, but from experimental data. Numbers of neuron pairs for within- and across-hemisphere lottery-lottery neurons are 6337 and 5959 respectively; for lottery-surebet neurons are 2641 and 2954 respectively; for contra-contra neurons are 44 and 39 pairs respectively. Note that mixed selectivity neurons are excluded from the last analysis. Vertical ticks indicate the median and 90% confidence intervals for each distribution.

With this architecture, we found parameters that qualitatively reproduce our optogenetic findings: namely, bilateral inhibition during either epoch weakened the relationship between ΔEV and lottery choices (Fig. 3c, Fig. 6d), and unilateral inhibition during the spatial planning but not abstract choice epoch resulted in a contralateral impairment and ipsilateral bias (Fig. 5b, Fig. 6d). Activity patterns in the resulting model units paralleled the electrophysiology data (Fig. 4), showing early-onset offer value and abstract choice signals that persisted into subsequent decision stages, followed by spatial action signals after offer locations were revealed (Fig. 6b,c). These results indicate that a network with dynamically shifting interhemispheric interactions is sufficient to recapitulate our causal and physiological results. To be clear, this mode switch from cooperative to competitive across epochs is not a result of changing weights. During the first epoch, strong mutual inhibition with balanced inputs keep the Contra-Contra subnetwork in a stable low activity state, which masks the interhemispheric competition. The whisker input drives the mode switch by providing a strong input to one hemisphere, breaking the symmetry while providing additional excitation to generate a bistable configuration of the Contra-Contra neurons.

To further validate this proposed mode switch, we examined noise correlations between simulated neuron pairs to characterize functional connectivity between distinct neural populations in the model. The model generated three key predictions, all empirically confirmed using simultaneous bilateral ALM recordings during task performance (Fig. 6f). First, consistent with bistable attractor dynamics for abstract decisions, neuron pairs with the same tuning (lottery-lottery) showed positively shifted noise correlations, whereas oppositely tuned neuron pairs (lottery-surebet) showed negatively shifted noise correlations (Fig. 6e,g left and middle). Second, in line with interhemispheric cooperation for abstract decisions, noise correlation patterns for abstract choice neurons were similar within and across hemispheres. Third, reflecting interhemispheric competition during spatial action selection, contralateral action-coding neurons from opposite hemispheres exhibited negatively shifted noise correlations (Fig. 6e,g right). We restricted the last analysis to neurons with pure spatial tuning, serving as a proxy for output populations driving contralateral movement. Taken together, these simulations and bilateral recordings suggest that frontal motor cortical hemispheres operate as a distributed attractor network to maintain abstract decisions, which then transition into a competitive regime to implement spatial action selection.

## Discussion

Using a novel economic decision task in mice (Fig. 1), we reveal a neural mechanism that transforms abstract offer values and economic choices into spatial actions across temporally dissociable decision stages. Mesoscale imaging and causal survey of the dorsal cortex identify the frontal motor cortex, in particular the anterolateral motor cortex (ALM), as a critical locus for both computations (Fig. 2, Fig. 3). During the first epoch, ALM neurons encode option values and choices before offer locations are revealed (Fig. 4), and silencing ALM activity produces non-lateralized impairments in economic decision preference (Fig. 3, Fig. 5). These findings support a model in which values are compared in the abstract space of options independent of sensorimotor contingencies^10^. Such computations may be instantiated via mutual inhibition between abstract “goods” attractors^21,40,41^, distributed across both hemispheres (Fig. 6). After offer locations are revealed, multiplexed ALM neurons integrate value and spatial information to guide action selection (Fig. 4), similar to activity reported in primate lateral prefrontal cortex^20,42^. Furthermore, we show that these signals are causal to task performance, and the eventual spatial choice may rely on competitive ALM dynamics between hemispheres (Fig. 5, Fig. 6).

Although we attributed the neural responses and causal contributions in the first epoch to non-spatial, value-guided economic choices between lottery versus surebet offers, an alternative interpretation is that these signals represent opposing sensorimotor rules (lick towards or away from the upcoming whisker stimulus). If mice were solving the task using a lick-toward versus lick-away rule, then mice with opposite contingency mappings (whisker stimulation signals lottery location vs whisker signals surebet location) would show opposite behavioural and neural signatures for lottery and surebet choices. However, unlike previous studies of rule-based behaviours^43^, we did not observe such phenomena (Fig. S2). Instead, we found that mice from both sensorimotor rule mappings were comparatively faster for lottery choices compared to surebet choices (Fig. S2e); and widefield imaging revealed higher cortex-wide activation for lottery choices compared to surebet choices, irrespective of sensorimotor rules (Fig. S5d). Together, this suggests that in our task, abstract value-guided choice is a more parsimonious description for animals’ strategies than “lick toward versus lick away”. Finally, it is important to point out that although we cannot definitively eliminate the sensorimotor rule hypothesis, that would not change the central conclusion in our study. Indeed, the mechanism of dynamic circuit reconfiguration from distributed, non-spatial computation to lateralized motor processes may generalize to other forms of decision-making, including context-based perceptual decision-making and rule-guided behaviours^44–48^. Future extensions of the task to include multi-modal value cues and multi-effector motor reports beyond two-alternative forced choice lick responses may help differentiate these more nuanced explanations.

Consistent with other mesoscale imaging studies^35,37,49^, we observed distributed representation of task information across the dorsal cortex, even when the underlying computation is isolated from spatial action preparation and execution (Fig. 2). These widespread dynamics may reflect correlated fluctuations in arousal, reward expectation, or embodied behavioural strategies, alongside neural representations of cognitive variables essential for task performance^35,50,51^. By contrast, the corresponding causal map for choice, revealed by our optogenetic survey, is not widespread but specific to the frontal motor circuit (Fig. 3)^30,52^. This causal map is also considerably more restricted compared to those found in decision-making studies using virtual reality paradigms^37^, which had more enriched task environments that may recruit broader cortical participation. Complementary cellular resolution recordings further reveal how non-linear transformations from abstract decisions to spatial actions can be implemented at the single-neuron level (Fig. 4). Although our findings identify ALM as a key node for transforming value-guided decision to spatial actions, our investigation is confined to the dorsal cortex. It remains to be tested whether these computations are locally instantiated in ALM or arise through recurrent interactions between ALM and other deep cortical and subcortical loops^22,53–55^.

The observed mode switch in interhemispheric interactions from cooperation to competition across task epochs is reminiscent of findings that early but not late unilateral ALM inhibition can be compensated by the unperturbed opposite hemisphere^56^. Hemispheric interactions may also be asymmetric, with the direction of influence dependent on task-related hemispheric dominance^57^. Here, we propose a computation-specific organization of frontal motor subnetworks^58^, where abstract choice neurons in both hemispheres form a distributed attractor network; and spatial action neurons encoding the contralateral motor command have competitive interactions between hemispheres (Fig. 6). How these ALM subnetworks map onto different cortical layers (Fig. S9d,e), projection targets^31^ and molecularly-defined cell types^59^ remain to be further examined to constrain the underlying microcircuit logic. Another open question is whether decisions are formed through sequential bistable attractor dynamics^60^, first in an abstract value space and later in an action space, or by a context-dependent conjunctive attractor network^61,62^. Exploring alternative dynamical model architectures could generate testable experimental predictions to differentiate these hypotheses.

Previous studies have proposed that abstract representation of value, as identified in primate orbitofrontal cortex (OFC)^13^, may have emerged late in evolution alongside the expansion of the frontal lobe^10^, since value signals observed in rodent OFC appeared spatially selective^22^. Leveraging a behavioural task conceptually aligned with primate paradigms of economic decision-making^20^, our study demonstrates that abstract value representation can be present in rodents when these computations are isolated from spatial processes, providing additional support for frontal motor contributions to non-spatial cognitive processes^63,64^. Together, our findings illustrate how dynamic circuit reconfiguration enables the transformation of cognitive decisions into concrete actions, providing a general principle for flexible control of adaptive behaviour.

## Methods

### Animals and surgical procedures

All experiments were performed in line with the UK Animals (Scientific Procedures) Act of 1986 (PPL: PD867676F) after local ethical approval by the Sainsbury Wellcome Centre Animal Welfare Ethical Review Body. A total of 34 adult mice (20 males, 14 females) at least 8 weeks old at the start of experiments were used in this study. Nine mice (CaMKII-tTa tetO-GCaMP6s) were used for task development and provided behavioural data. Six mice (CaMKII-tTa tetO-GCaMP6s) mice were used for widefield calcium imaging experiments. Eight mice (VGAT-ChR2-EYFP) were used for optogenetic inactivation experiments. Six mice (CaMKII-tTa tetO-GCaMP6s, including 1 mouse used in widefield imaging experiments) were used for optogenetic control experiments. Four C57BL/6J mice were used for electrophysiological recordings (2 for unilateral recordings, 2 for bilateral recordings). Two mice (VGAT-ChR2-EYFP) were used in electrophysiological validation experiments confirming optogenetic silencing.

For headplate implantation surgery, mice were anesthetized with isoflurane (5% induction, 1.5% maintenance) and injected subcutaneously with an analgesic (Metacam). The scalp was shaved and cleaned with an antimicrobial cleanser (Hibiscrub) and sterile saline. Mice were fixed in a cranial stereotaxic frame (Kopf Instruments) and placed on a heat mat (37°C). Body temperature was monitored via a rectal probe, and lubricant was applied to the eyes. Scalp was removed to expose the dorsal surface of the skull, which was cleaned and dried. The remaining scalp was adhered to the skull using tissue adhesive (Vetbond; 3M). A metal headplate was fastened along posterior edge of the skull with clear dental cement (Super-Bond C&B, Sun Medical) allowing access to the dorsal surface of the cortex. The exposed skull was then covered with a thin layer of dental cement. Two locations on the skull were labelled using a marker pen (bregma and +3 mm anterior from bregma) to aid stereotaxic alignment in later experiments. For electrophysiological access to the brain, a small targeted craniotomy was performed 1-2 days before recording experiments (Volvere i7 E-Type Micromotor Drill System; NSK). All surgeries were performed using aseptic techniques.

After headplate implantation surgery, mice were single-housed in individually ventilated cages equipped with environmental enrichment. Cages were maintained at ambient temperature and humidity and 12 h reversed day-night cycles. Mice received post-operative analgesia (Metacam) for a minimum of 2 days. Following recovery (~ 1 week), mice were water-restricted prior to the start of behavioural training, with *ad libitum* access to food. Mice received water (~ 1 ml per day) through daily behavioural training. Supplementary water was provided if necessary for mice to maintain a stable body weight (80 - 85% of their starting weight). On non-training days, 1 ml of water was provided each day.

### Behaviour training

#### Hardware

Head-fixed behavioural training was conducted in custom-made aluminium training boxes (50 × 50 × 50 cm) lined with sound-attenuating foam. Head fixation was achieved using two clampable metal arms equipped with screws that secured the headplate from both sides. During training sessions, mice were positioned inside a perspex tube that supported the body while the head was fixed. Auditory stimuli were delivered via an electrostatic speaker (ED1 speaker with ES1 driver; Tucker-Davis Technologies) and calibrated using a microphone (70 dB, UMIK-1; minidsp microphone; REW software; www.roomeqwizard.com). For controlled stimulation of the whiskers, two compressed air lines (calibrated at 2 bar pressure; R412006111; Emerson-Aventics) were directed to the underside of the left and right whiskers using air-pipe tubing (3 mm diameter; PUN-3X0,5-SI; Festo) gated by 2 solenoid valves (EV-2-12; Clippard). Lickspouts were pieces of 10 cm long 1/16” x 1/32” steel tube (Coopers Needle Works), and were gravity-fed water via rubber tubing gated by two solenoid pinch valves (225P011-11; NResearch). Unitary reward volume (1 µL) was calibrated using high-precision weighing scales and standardised across left/right reward lines and across training rigs. Target reward volumes were delivered as a train of unitary rewards at 10 Hz (e.g. 3 µL would be a sequence of 3 unitary 1 µL rewards). Air and water solenoid valves were mounted on the exterior of the behaviour chamber. The lickspouts were fixed in a custom printed holder coupled to two servo-motors (XL-320; Robotis), enabling translations in the forward/backwards and left/right axes. Forward/backward translation of the lickspouts signalled the onset/offset of the response window. Left/right position adjustments of the spouts were enabled in closed-loop fashion during training and pre-training to correct for spatial choice biases (adjustments in ± 0.5 mm steps). Our lick detection system is based on a design developed at Janelia Research Campus (https://github.com/janelia-experimental-technology/ Comparator-Dual-Lick-Detector). We also recorded videography via two infrared-sensitive cameras (Raspberry Pi High Quality Camera; SC1220; with a C-Mount 8 − 50 mm Zoom lens; WAV-18245; The PiHut) positioned face and side on to the animal (Fig. S4a-b). The behaviour chamber was illuminated with an infrared LED array and an ambient white light, to facilitate recordings of pupil responses. We recorded video at 30 frames-per-second (fps) during training and 90 fps during experimental sessions. Data were acquired with Raspberry Pis (RPi4B running camera control software https://github.com/larsrollik/rpi_camera_colony, 1 for each camera) which also recorded timestamped TTL signals to synchronise frames to trial (lottery sound cue) onset.

Behavioural hardware and task design was controlled by a modified version of Bpod (https://github.com/sanworks/Bpod_Gen2) in Matlab, with additional in-house software and custom-made printed circuit boards (PCBs) to support training automation, power management, auditory stimulus delivery, lickport motor actuation and lick detection. Hardware and software designs are freely available at https://gitlab.com/sainsbury-wellcome-centre/delab/bpod-auto.

#### Task design

Head-fixed mice were trained on an economic choice task designed to temporally separate abstract value-guided decision-making and spatial motor planning processes. On each trial, mice were presented with a choice between a guaranteed water reward (Surebet; 3 µL; P(reward) = 1) or a probabilistic water reward (Lottery; 0 to 16 µL; P(reward) = 0.7). Mice reported their choice with directional licking. Across trials, both the lottery reward size and location (left or right) were randomised and cued by two sequentially presented stimuli. First, a transient auditory stimulus (Value cue; 1 s duration) signalled the current lottery offer. Twenty-eight mice were trained with pure tone frequency discrimination (4.6, 7.4, 9.8, 14, 20 kHz), and 4 mice were trained with click rate discrimination (click rates = 10, 45, 60, 81, 110 Hz, individual clicks were 9 kHz short pure tones each lasting 3 ms). Following the value cue, a lateralised air-puff stimulated either the left or right whiskers (Location cue, 10 Hz pulse train, 50 ms pulses; 1 s duration), signalling the spatial mapping of lottery and surebet offers onto left and right lick spouts. This means that when presented with the same offer pair, on some trials mice had to lick left and on others lick right to report the same offer choice. Moreover, the sequential delivery of value and location cues means that during the first cue epoch, information is available to guide a non-spatial ‘lottery vs surebet’ choice, referred to as abstract choice; but only after the onset of the second cue can mice plan a specific directional licking response to enact their choice. After the cue period (0 − 2 s), a motorised dual-lick spout extended forwards, providing a salient audiovisual ‘go’ cue, which allowed mice to report a choice. Full extension took ~140 ms (total distance 1 cm, Fig. S4a,c). The response window was 3 s, if no choice was reported, the trial was classified as a miss trial. If a choice was made (defined as the direction of the first lick contact detected on either spout), a further 4 s was allocated to allow potential reward consumption before the lickport retracted back into the starting position for the next trial. Response time (RT) was defined as the time to the first lick after the onset of the response window. A variable inter-trial interval lasted between 3 to 5 s. Sessions were terminated by the experimenter following satiation (usually ~ 5 consecutive miss trials).

We counterbalanced stimulus contingencies across mice (Fig. S2). Some mice learned a positive auditory frequency-to-value mapping (i.e. high frequency, large lottery payout) and others learned a negative map (i.e. high frequency, small lottery payout). We also counterbalanced the whisker-to-offer mapping contingency across mice. For some mice, the side of the whisker stimulus cued the lottery location (i.e. whisker left, lottery left); and for others, the whisker stimulus cued the surebet location (i.e. whisker left, lottery right).

#### Pre-training and habituation

Water-restricted mice were first habituated to handling and head-fixation (1 − 2 days). While head-fixed, mice were encouraged to drink water offered via a handheld syringe. Once acclimated to head restraint, mice spent 3 - 5 days in a pre-training routine designed to teach proactive left/right licking for reward. In the first phase, the lick spouts were positioned close to the mouth and droplets of water (3 µL) were triggered by lick events (inter-reward interval 0 − 1 s). If mice did not lick for 20 s, a left/right reward was randomly delivered. Mice were free to lick either side for reward. However, if a lateralised bias was detected (≥ 80% bias across 20 consecutive rewards), the lickport position updated in a closed-loop fashion to counteract this bias. After each left or right motor adjustment (1 ‘step’ = 0.5 mm), further adjustments were disabled for 20 additional trials, after which the bias was re-assessed. While there was no limit on the number of lickport adjustments that could be made in the session, a maximum offset from the starting position of 3 corrective steps was enforced (i.e. ± 1.5 mm from centre). The next phase of pretraining introduced forward/backward lickport movement. On each trial, the lickport extended (for 5 s) and then retracted, requiring mice to wait between rewards (inter-trial interval 0 − 2 s). Rewards were also now structured in interleaved, side-specific 3-trial blocks. This encouraged mice to alternate left/right choices in order to keep receiving rewards. Incorrect licks were not penalised. Rewarded side switched after 3 rewards were successfully collected. Lickport position adjustments were also enabled, shifting after 3 consecutive incorrect trials. In practice, reward omission on one side usually prompted exploratory licks to the opposite side, so few motor adjustments were typically required.

#### Task training

##### Training overview

Mice learned the economic choice task via a series of training stages designed to shape performance through operant conditioning (Fig. S1a; Training: Stages 1 - 3; Final task: Stage 4). Optimal performance in the full task required learning two distinct associations: (1) the mapping between auditory stimuli and offer values/outcomes and (2) the spatial mapping between lateralised whisker stimulation and offer locations. To simplify the learning process, the training regime prioritised acquisition of the auditory-value mapping with more gradual learning of the flexible spatial response contingency. For the majority of training, we restricted the trial-set to two lottery offers cued by high (20 kHz) vs low (4.6 kHz) auditory tones respectively: a bad lottery offer (0 µL) and a good lottery offer (16 µL). Accordingly, mice should develop clear value-guided choice preferences across trial-types (e.g. choosing the 3 µL surebet on 0 µL lottery trials while favouring the lottery on 16 µL lottery trials). During Stage 1, offer locations were fixed across trials and training sessions, e.g. the lottery was always presented on the left lick spout and surebet on the right (starting configuration was counterbalanced across mice). At this stage, value-guided performance can be described as a mapping between auditory stimulus and lick direction. However, once choice behaviour met a performance threshold (ΔP(Lottery) ≥ 0.7 across the two trial-types), the location mapping was reversed (i.e., if the lottery was previously on the left, it would now switch to the right and vice versa for the surebet). This change would be explicitly signalled, as the side of the whisker stimulus would also switch accordingly. Following a reversal, performance decreased due to perseveration of the initial learned contingency/strategy, but gradually recovered over subsequent sessions (Fig. S1b). Mice were retrained on the new offer location contingency until reaching performance criterion after which the offer locations were reversed again. With repeated exposure to performance-contingent reversals, mice required fewer sessions to recover performance (Fig. S1b-d). This indicates an increase in behavioral flexibility and learning of the instructive nature of the whisker stimulus. When mice were able to recover performance within a single training session (after 5-15 reversals, depending on learning speed across mice), we introduced within-session switches in location - initially in alternating reversal blocks (Stage 2), and then fully randomly interleaved (Stage 3). To maximize rewards during Stage 3, mice have to rely on the lateralized whisker stimulus to signal the offer-locations on each independent trial. Following stable performance on Stage 3, we introduced the intermediate lottery trial-types (Stage 4; Full psychometric task). On average, it took 66 ± 27.04 training sessions (mean ± std) for mice to reach the final stage of the task (3-4 months given 5 training sessions per week). Full details of each training stage are outlined below.

**Stage 1**. During the first few training sessions we presented low and high lottery trial types in alternating blocks (6–10 trials; Fig. S1e(i)). Once mice reached criterion discrimination performance (see below), the lottery trial sequence was randomized for the remainder of training (Fig. S1e(ii)). We evaluated value-guided discrimination as the difference in lottery choice probability between high and low lottery trial types (ΔP(Lottery)), measured within a 100-trial rolling window. For example, if P(Lottery) = 0.1 on low-lottery trials and 0.9 on high-lottery trials, the discrimination score would be 0.9 − 0.1 = 0.8. The maximum value across the rolling-window was taken as the session’s *peak discrimination score*. A peak score of ≥ 0.7 was used as the threshold for reversal. To avoid mice remaining in a single stage for too long, we also allowed for more lenient thresholds: two consecutive sessions with a peak score of ≥ 0.6 or three consecutive sessions ≥0.5. If mice met the threshold within a session (Fig. S1e(ii)), offer locations were reversed on the following day (Fig. S1e(iii)) and mice were retrained until performance recovered and they met the threshold for another reversal (Fig. S1e(iv)). During the early stages of training, the lottery offer had a high payout probability (P(reward) = 0.9). We gradually decreased this probability across reversals to reach the final P(reward) value of 0.7 across the first five reversals: [0.9, 0.85, 0.8, 0.75, 0.7]). The ‘high’ lottery reward magnitude increased across reversals to keep expected value approximately constant (high lottery reward across the first 5 reversals: [9, 10, 12, 14, 16 µL]). To promote associative learning between stimuli and reward outcomes, the auditory and whisker cues were initially presented in an overlapping configuration during Stages 1 and 2. The auditory stimulus lasted 2.5 s and the whisker stimulus lasted 1.5 s. The whisker stimulus was delivered at *t* = 1 s relative to auditory cue onset. The lickport extended at *t* = 2 s, meaning both auditory and whisker cues extended into the start of the response window.

**Stage 2**. The next phase of training involved within-session location reversals. Offer locations were reversed across alternating blocks of trials - initially in long blocks (40 - 60 trials; Fig. S1e(v)), and when the discrimination threshold was met, in short blocks (10 - 20 trials; Fig. S1e(vi)). As in Stage 1, location reversals were signalled by switches in whisker stimulation side.

**Stage 3**. Following criterion performance in Stage 2 (short block reversals), we fully randomized the offer locations across trials (Fig. S1e(vii)). During this stage, we also gradually removed the temporal overlap between value and location cues on a subset of trials. On these ‘no overlap’ trials, the value cue (1 s) and location cue (1 s) were presented sequentially, as in the final version of the task (Fig. 1c). To facilitate this transition, we randomly interleaved ‘no overlap’ trials with the ‘overlapping cue’ trials, increasing the relative proportion of ‘no overlap’ trials across training sessions in a performance-dependent manner.

**Stage 4**. Once mice reached the criterion performance in Stage 3 sessions with 100% ‘no overlap’ cue trials, we introduced the remaining intermediate lottery offer trials (Fig. S1e(viii)). We initially introduced the intermediate trials with low probability, increasing the proportion of across sessions until all five lottery trial types ([0, 2, 4, 8, 16 µL]), and both offer location trial types, were fully randomized.

Across stages 1-3, we implemented two shaping features in closed-loop fashion to correct response biases: (1) spatial bias correction and (2) abstract bias correction. First, as described above, we dynamically adjusted the lickport position if spatial bias was detected. We monitored spatial choices across a 50-trial rolling window, and implemented a lateral shift (0.5 mm) in lickport position if a bias (≥ 80 %) was detected. If a shift was made, further adjustments were disabled for the subsequent 50 trials. A maximum offset of 1.5 mm (3 steps) from starting position was allowed. Second, we implemented ‘forced trials’ to instruct appropriate offer choices with respect to ΔEV trial type. For example, if mice continually chose the lottery indiscriminately (bias ≥ 80 % assessed across a 20-trial rolling window), ‘forced trials’ would be transiently enabled until the bias was corrected (P(Forced trial) = 0.9). For instance, during a period of lottery bias, only the surebet reward was available on a forced trial. This means that licks at the ‘lottery’ spout would be ignored, and the surebet reward would be delivered if mice subsequently licked the surebet spout at any point in the trial. If no response was made in 30 s, the trial was aborted and classed as a miss. The inverse would happen if surebet choice bias was detected. This aimed to steer mice away from a habitual strategy of always choosing lottery or surebet, and/or forming a directional response conditioned on the whisker stimulation side. Spatial and abstract bias corrective procedures were disabled during all recording and optogenetics experiments.

### Behavioural performance

#### Psychometric fits

To characterise lottery choice preference across offers, we fit a logistic regression model to mice’s binomial choice data, where the lottery choice was fit as a logistic function of ΔEV (*EV*_*lottery*_ − *EV*_*surebet*_) and bias (excluding miss trials):

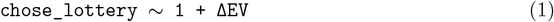

Psychometric curves were generated by calculating model-predicted lottery choice probability across a continuous range of simulated ΔEV values, and displayed with the observed data points for each unique ΔEV trial type (mean ± 95% binomial confidence intervals).

#### GLM-HMM fits for behavioural and widefield analyses

To examine the relationship between task performance and behavioural state, we fit a hidden Markov model (HMM) with a Bernoulli GLM (GLM-HMM)^33,34^ to psychometric data from each animal. The model comprised: 1) a transition matrix specifying the probability of transitioning within and between latent states across trials, and 2) a set of GLM weights describing choice behaviour within each state as a function of five input regressors: current trial ΔEV, overall lottery bias, overall spatial bias (encoded as whether the lottery location was on the left), perseveration of lottery choice from the last trial, and perseveration of spatial choice from the last trial (encoded as whether lottery choice on the current trial would repeat the spatial choice of the previous trial). This framework allowed us to identify subject-specific latent behavioural states characterised by varying levels of stimulus-dependent task engagement, choice biases (value and/or spatial), and trial-history dependent strategies. We applied a threshold to the posterior state probabilities estimated by the model, and only trials where P(State) ≥ 0.7 were assigned the corresponding state.

We compared performance of different models with varying numbers of latent states (2-5 states) using 5-fold cross-validation across sessions^33^. On average, model predictive performance saturated at around 3 states, and incorporating more than 3 states resulted in marginal improvements in the model fit to the data (Fig. S3a). Moreover, the proportion of trials assigned to the 4th and 5th states were comparatively low (<10%), and 4- and 5-state models had increased unclassified trials (P(State) < 0.7) compared with the 3-state model (Fig. S3b). Therefore, we opted for the 3-state GLM-HMM model for the rest of the study.

We defined a task-engaged state as the state with the greatest positive weight for ΔEV. This state usually had minimal weights for the other regressors indicating that lottery choices in this state were primarily guided by lottery offer value. The remaining two states, usually well described by leftward vs rightward choice biases (Fig. S3c,f), were combined into a singular ‘disengaged’ state. We separately computed psychometric performance during engaged and disengaged trials as described above. Compared with spatial choice bias, abstract choice bias did not appear to be as strong a feature of disengaged state performance (Fig. S3g), suggesting that habitually choosing the lottery or surebet offer were not common behavioural strategies. However, mice did show a small surebet choice bias in at least 1 of the two disengaged states (Fig. S3g).

#### Rolling window trial inclusion for optogenetic experiments

Because animals’ baseline behavioural states also fluctuated during the optogenetic perturbation sessions, we aimed to identify task-engaged state on control trials in these sessions and assess how cortical perturbations affected behaviours while animals performed the task in a stimulus-dependent manner. However, since laser ON trials that targetted 17 different cortical regions were randomly interleaved with laser OFF control trials, GLM-HMM that fit state transitions across all trials was not suitable for optogenetic sessions. Instead, we used a simpler, model-free method and tracked discrimination (ΔP(Lottery) between high-lottery and low-lottery trials) on laser OFF control trials in a 100-trial rolling window, and set an inclusion criteria of discrimination ≥ 0.7. If discrimination within the 100-trial window met this threshold, all trials within the set were included for analyses (including the interleaved laser ON trials). The rolling window was performed in the forward direction, meaning that if trial 1-100 met the threshold, these trials were banked as ‘good’ trials, but trials 2-101 failed to meet this threshold, only trial 101 would be labelled as ‘bad’ - with the assumption that the additional of the new trial (trial 101) now caused the drop in discrimination. If performance recovered again later in the session (e.g. if trials 3-102 met threshold again), trial 102 would be included. While this method was not as systematic as GLM-HMM, it avoided potential confounds of fitting transitions across interleaved laser ON trials belonging to different optogenetic experiments, and allowed us to identify blocks of control trials with good task performance and include laser ON trials within these blocks for analyses.

#### Response time

We define response time (RT) as the time of the first detected lick at either lick spout relative to the lickport extension onset time and report median RTs calculated across trials. When reporting changes in RT following optogenetic manipulation (Fig. 3, Fig. 5, Fig. S7), we calculated the difference between laser ON RTs and the median of the session-matched laser OFF trial RTs. As the response window onset was defined by lickport extension on each trial, we also used videography to assess anticipatory licking outside of the response window (Fig. S4d-e). These analyses confirmed that mice do not lick prematurely during the value and location cue epoch, and most lick responses occurred within the designated response window.

### Videography

#### Pre-processing and analysis

To incorporate trial-evoked movements into our widefield imaging analyses, we used SLEAP^65^ to track orofacial and body movements, and pupillometry, in the behavioural videography data (Fig. S4). We trained a subject-specific model on human-labelled frames selected at random across sessions (typically ~ 100 frames per model), with additional frames added to the training set if tracking quality was poor. We trained separate models to track pupil (side camera) and forepaw nodes (front camera), and a third model (side camera) to collectively track tongue, jaw and nose nodes. We also tracked lickport position as a secondary method to check video alignment to trial events. To quantify licking, we tracked the tongue tip across video frames and detected frames when the tongue was present by thresholding the tongue node confidence score (threshold = 0.1, Fig. S4d-e). To quantify pupil size, we tracked 8 points outlining the edge of the pupil and extracted the area of an ellipse fit across the points (Fig. S4a right). For each trial, the pupil trace was z-scored using a pre-stimulus baseline (−1 to 0 s relative to sound cue onset; Fig. S4f). To quantify nose (Fig. S4g), jaw (Fig. S4h) and paw (Fig. S4i) movements, we computed the euclidean displacement of each tracked node across consecutive frames and converted this to velocity (mm/s). We also computed the euclidean distance relative to the mean position during the pre-stimulus baseline. To quantify whisking motion we outlined a rectangular region of interest (ROI) over the left and right whisker pads (front camera, Fig. S4b) and calculated frame-to-frame correlation of the vectorized pixel values. We defined whisking motion as 1 minus the frame-to-frame correlation such that periods of quiescence (which would have high frame-to-frame correlation) would have low motion scores and vice versa (Fig. S4j). All video traces were smoothed with a causal moving window of 100 ms (9 frames at 90 fps). For inclusion in widefield imaging analyses, 90 fps videography traces were downsampled to match the imaging sampling rate (30 fps).

### Widefield imaging

#### Acquisition and hardware

Imaging was performed on a tandem-lens widefield fluorescence macroscope (85 mm f/1.8D objective, 50 mm f/1.4D tube lens, Nikon; described previously^49^). Images were captured on a CMOS camera (PCO.edge 5.5) via an emission bandpass filter (5256/50-25; Semrock). Frames were recorded at 60 frames per second (fps), with interleaved blue (470 nm; Thorlabs LED; M470L4, with excitation filter FF02-447/60-25, Semrock) and violet (405 nm; Thorlabs LED; M405L4, with excitation filter FF01-405/10-25, Semrock) excitation combined via a dichroic (FF458-Di02-25 × 36, Semrock). A second dichroic (FF495-Di03, Semrock) directed excitation down to the sample while allowing transmission of emitted green fluorescence up to the camera. Alternating blue/violet LED excitation was controlled via a Teensy microcontroller and data acquisition card (USB-6001; National Instruments)^66^ and synchronised with the camera rolling shutter acquisition trigger output. Acquisition control was provided via Matlab-based software (‘WidefieldImager’; https://github.com/musall/WidefieldImager). We acquired 9 s of imaging data on each trial; 3 s before and 6 s after onset of the value sound cue.

#### Data pre-processing

Widefield imaging data were pre-processed using existing software^35,66^. Briefly, to reduce computational cost, we restricted our analyses to a truncated window 1 s before to 5 s after sound cue onset. To compress the data, we then used singular value decomposition (SVD) to compute the 2000 highest-variance dimensions across the time-series for each recording session, yielding spatial components (pixels x components) and temporal components (components x time points). Temporal components were high-pass filtered above 0.01 Hz using a zero-phase, second-order butterworth filter. Calcium dependent fluorescence signals were isolated from intrinsic signals using hemodynamic correction across blue and violet excitation channels. Following hemodynamic correction, effective imaging sample rate was 30 fps. For each trial, Δ*F/F*_0_ was then computed using the mean fluorescence during the 1 s pre-cue baseline as *F*_0_. Following pre-processing, spatial and temporal components were multiplied to recover data in original pixel space.

For each animal, pre-processed data from each session were then rigidly aligned using landmarks based on macro vasculature. Session-concatenated data were then aligned to the Allen Common Coordinate framework (CCF) using four anatomical landmarks (left, centre and right points where frontal motor cortex meets the olfactory bulbs, and the base of retrosplenial cortex) and two functionally-derived landmarks (stimulus-triggered average (STA) images following left vs right whisker stimulation revealing barrel cortex). All subsequent analyses were performed on CCF-aligned session-combined imaging data. The full dataset comprised 6 CaMKII-tTa tetO-GCaMP6s mice; 52 experimental sessions; 8.67 ± 2.01 sessions per mouse; 2156.8 ± 586.32 trials per mouse; 1300.7 ± 334.6 engaged trials per mouse; 400.12 ± 145 disengaged trials per mouse (mean ± std).

### Widefield imaging analyses

#### Encoding model

To examine dorsal cortex representation of task information, we analysed CCF-aligned widefield imaging data for each animal using a linear regression model^35^. To separate contributions of task and movement related signals, the model design matrix included stimulus and behavioural event regressors and videography-based regressors that captured motor behaviour. An overview of inputs to the model is provided in Tab. 1. Event regressors were time-expanded to capture temporal relationships between task regressors and cortical fluorescence. For example, for each lottery-value trial-type, the design matrix contained multiple time-shifted copies of a binary event pulse, forming a block of regressors spanning 0.5 to +4 s relative to sound-cue onset. This ensured that regressor kernels included a short pre-stimulus baseline and extended into the response window. Similarly, whisker stimulus side, abstract choice and spatial choice regressors were also time-shifted across the same − 0.5 to 4 s window. We also included time-shifted regressors encoding reward pulse delivery times, and absolute time in the trial. In addition to the above task event regressors, the design matrix also included analog regressors derived from behavioural videography. These included whisking, nose, jaw, paw, pupil and tongue movement velocity. We opted for key-point based feature tracking, rather than including video components derived from dimensionality reduction methods^35,51^, for ease of concatenating design matrices across different recording sessions. For all videography-based analog regressors we also computed corresponding event kernels by binarising each trace according to a z-score threshold. Using a rolling baseline of 200 ms, we binarised each analog regressor according to a z-score threshold of 3 above baseline. Following an event, we ignored repeated events for an additional 100 ms. Movement-based event regressors were time-shifted using a − 2 to 2 s window to capture preparatory movement related activity. The model was fit using ridge regression, with the regularization penalty *λ* estimated independently for each pixel time-series in the widefield data via marginal maximum likelihood estimation^35^. Model performance was assessed using 10-fold cross-validation across trials, where the model was trained on 9 folds and used to predict the held-out fold. The full predicted and observed data set were then compared pixel-wise, and the squared Pearson correlation was computed to yield *R*^2^ maps summarising the fraction of variance explained by the model Fig. 2d.

**Tab 1:**
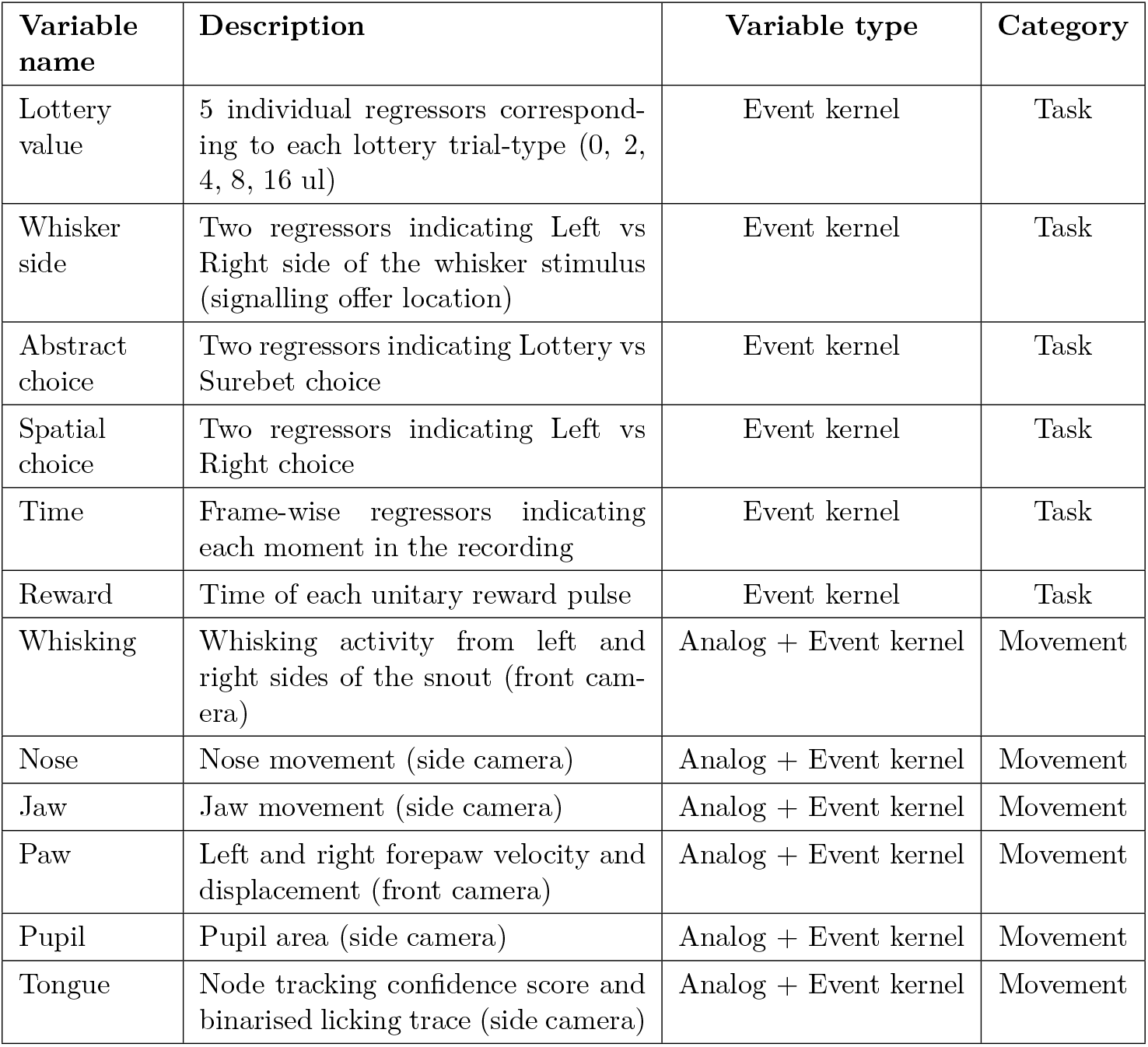
Summary of regressors included in the design matrix for widefield imaging analysis.

#### SVM Decoding

To assess the amount of task-related information contained in widefield signals, we trained separate linear support vector machine (SVM) classifiers (L2 regularised with *C* = 1) to decode the 1) whisker stimulus side (left vs right), 2) abstract choice (lottery vs surebet), and 3) spatial choice (left vs right) on each trial. Widefield activity was first pixel-averaged per the 59 regions defined by the Allen atlas, and concatenated across sessions for each mouse. We then divide the trials into those during the engaged state (*P* (engaged ≥ 0.7) from the GLM-HMM) and the disengaged states (*P* (disengaged ≥ 0.7) for either of the two disengaged states). Within each state, we randomly subsample trials such that there are equal number of trials for each combination of whisker stimulus and choice sides, such that there is no difference in distribution of whisker stimulation locations, spatial choices and abstract choices. After this procedure, we balance the number of trials in engaged and disengaged states such that differences in decoding performance between the states cannot be attributed to a difference in the amount of data available. Widefield activity were aligned from −1 to 4 s relative to sound onset, and we performed decoding on each 250 ms window with a step size of 33.33 ms (constrained by the 30 Hz sampling rate of widefield imaging). We performed 2-fold cross validation with 10 repeats and reported the mean accuracy. To obtain the null distribution of decoding accuracy, we shuffled the identity of the decoded target across trials followed by the same decoding procedure and report the mean across 50 shuffles. Finally, we calculated the mean decoding accuracy across all mice (*n* = 6) along with the standard error around the mean.

For single brain region decoding, we repeated the same decoding procedure but instead used the activity of a single region as input for each decoder. For cross-window decoding, we also used the same training procedure, but instead tested the decoder on widefield activity from a different time window and again reported the mean accuracy of the decoder.

### Laser scanning photoinhibition

#### Hardware

We performed laser-scanning optogenetic inactivation using a blue laser coupled to a galvo-scanning system (Zapit^36^; https://github.com/Zapit-Optostim/zapit). Briefly, 473 nm laser light (OBIS LX 75 mW, Coherent) was directed into a pair of dual-axis galvometer scan mirrors (ScannerMax Saturn 5B; Edmund Optics 16-040), then deflected downwards onto the skull via a dichroic mirror (Thorlabs; MD498). An objective (f = 100 mm, composed of two Thorlabs AC254-200-A achromatic doublets arranged in Plössl configuration) focused the beam onto the sample while also allowing light emitted from the sample plane to pass up to a camera (Basler ace acA1920-40 µm Monochrome USB 3.0; Edmund Optics 33-980) via a tube lens (f = 50 mm, composed of two Thorlabs AC254-100-A achromatic doublets in Plössl configuration) and emission filter (Chroma AT495LP). By comparing the resulting position(s) of the laser spot on the sample, imaged via the camera, to a set of pre-determined target points, scanner and sample reference frames were coregistered via an affine transformation. Following this calibration, laser light could be accurately targeted to any 2D location in the sample field of view. Subsequent registration of the sample to the Allen Common Coordinate Framework enabled specification of target location sites as stereotaxic coordinates relative to bregma (which was marked on the skull surface). Both scanner and sample calibration steps were semi-automated by the Zapit control software^36^. The software also allowed flexible control of laser stimulus parameters such as waveform design, timing and power. Reported power values are time-averaged and were measured at the sample plane (Thorlabs; PM100D2).

For bilateral inactivation experiments, the beam was rapidly scanned between paired target points across hemispheres at 40 Hz, with a travel time of 0.5 ms between locations. During non-stationary scanning periods, the beam was blanked to avoid between-site stimulation. For behavioural experiments we used square-wave pulses of 12.5 ms duration (duty cycle 50% per location). For electrophysiological calibration experiments, we used half-sinusoidal pulses to minimise photoelectric artefacts - matching time-averaged power of the laser stimulus to the square-wave laser stimulus used in behavioural experiments. For single-site (unilateral) stimulation, the beam was directed to the respective target and cycled on/off at 40 Hz with 50% duty cycle. The beam-path, optics and scan mirrors were housed in a custom machined enclosure to minimise mechanoacoustic cues. A blue LED (PiHut; FIT0878) fixed near the objective aperture provided a visual masking stimulus on photostimulation (laser ON) and control (laser OFF) trials. The masking light was triggered for 2 s spanning both the auditory and whisker cue periods using a continuous square wave. Laser stimulation onset was triggered using hardware triggers (TTL) aligned to sound cue onset, with highly reliable temporal precision and sub-millisecond latency^36^.

#### Experimental design

To identify cortical areas causal to task performance, we surveyed 17 bilateral regions spanning the dorsal cortical surface (stereotaxic target coordinates in mm relative to bregma): (2.5 A ± 0.5 L), (“ALM”: 2.5 A ± 1.75 L), (1.25 A ± 0.5 L), (1.25 A ± 1.75 L), (1.25 A ± 3 L), (0 A ± 0.5 L), (0 A ± 1.75 L), (0 A ± 3 L), (−1.45 A ± 0.5 L), (−1.25 A ± 1.75 L), (−1.25 A ± 3.5 L), (−2.7 A ± 0.5 L), (−2.5 A ± 1.75 L), (−2.5 A ± 3 L), (−2.5 A ± 4.25 L), (−3.75 A ± 1.75 L), and (−3.75 A ± 3 L). In some experiments, we also included unilateral ALM photoinhibition trials in the set of targeted sites (19 target locations in total). We randomised targeting across the different photoinhibition locations, and delivered photoinhibition on 50% of trials drawn at random across all behavioural conditions. The photoinhibition epoch (0 − 1 s, “abstract choice” epoch; or 1 − 2 s, “spatial planning” epoch) and laser power (2 or 4 mW) was also randomised across trials. Unless otherwise stated, we concatenated photoinhibition trials across laser powers for statistical power. For epoch-specific inactivation experiments (duration = 1 s) we delivered 800 ms fixed amplitude 40 Hz pulsed laser stimulation with 200 ms linear rampdown to attenuate electrophysiological rebound effects after laser offset.

We performed the same experiment in two cohorts of animals; (1) mice expressing channel-rhodopsin in inhibitory neurons (n = 8 VGAT-ChR2-EYFP mice) and (2) control mice that did not express an optogenetic actuator (n = 6 CaMKII-tTa tetO-GCaMP6s mice). Due to the large number of unique optogenetic conditions (38 conditions; 19 target locations x 2 epochs), we concatenated data across sessions and mice. We filtered out trials that did not meet task performance inclusion criteria and analysed data from VGAT-ChR2 and control mouse cohorts separately.

For experiments in VGAT-ChR2 mice, the data-set comprised 8 mice, 240 sessions, 18301 laser OFF control trials and 18675 laser ON trials. For bilateral photoinhibition during the abstract choice epoch (0 − 1 s), the dataset included 482.2 ± 9.5 trials per target region (mean ± std). For bilateral photoinhibition during the spatial planning epoch (1 − 2 s), the dataset included 488.7 ± 12.4 trials per target region (mean ± std). The unilateral ALM photoinhibition dataset included 382 left and 389 right hemisphere abstract choice epoch photoinhibition trials and 370 left and 361 right hemisphere spatial-planning epoch photoinhibition trials. We combined left and right hemisphere photoinhibition trials together to examine unilateral effects on behaviour.

For control mice, the data-set comprised 136 sessions, 12978 control trials and 13056 laser ON trials. For laser delivery during the abstract choice epoch (0 − 1 s) the dataset included 348.9 ± 11.4 trials per bilateral target region (mean ± std). For laser delivery during the spatial planning epoch (1 − 2 s), the dataset included 343.1 ± 11.6 trials per bilateral target region (mean ± std). The unilateral ALM dataset included 306 left and 319 right hemisphere abstract choice epoch laser ON trials and 337 left and 323 right hemisphere spatial-planning epoch laser ON trials.

For sub-epoch ALM photoinhibition experiments (Fig. 3d) we delivered 300 ms fixed amplitude 40 Hz pulsed laser stimulation with 200 ms linear rampdown (total duration = 500 ms). Photoinhibition (4 mW) was delivered randomly on 15% of trials at 0, 0.5, 1 or 1.5 s relative to the sound cue onset. The dataset comprised 73 sessions in 5 VGAT-ChR2 mice; 7446 control trials and 1247 photoinhibition trials; 311.8 ± 27.9 laser ON trials per sub-epoch.

#### Generalized Linear Mixed Models (GLMMs) of optogenetic effects

To examine putative epoch and site-specific effects of laser stimulation on task performance, we compared laser ON trials for each photoinhibition condition with the session-matched laser OFF trials using generalised linear mixed models (GLMMs). Specifically, to test whether laser stimulation had any effect on lottery choice behaviour, we compared predictive performance of a baseline model with a full model as follows (described using Wilkinson’s notation)^67^. The baseline model was

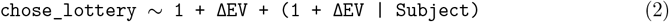

where the lottery choice on each trial was modelled as a function of an intercept and ΔEV (*EV*_*lottery*_ − *EV*_*surebet*_) on each trial as both fixed effects and random effects across subjects. The full model was

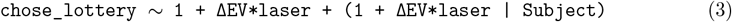

which incorporated an additional main effect laser, which indicated whether the laser was on or off on each trial, and an interaction term between ΔEV and laser. We performed 20-fold crossvalidation (CV) to evaluate model predictive performance on out-of-sample test data. If inclusion of the laser terms in the full model significantly improves model performance, this indicates that laser stimulation at that cortical site during that epoch has a measurable influence on lottery choice behaviour. We compared mean log-likelihoods across matched folds using a paired t-test, generating a *P*-value for each cortical target site and each epoch. Resulting *P*-values were corrected for multiple comparisons across cortical locations using the Bonferroni-Holm method (*α* = 0.05; 17 cortical locations). We performed the same procedure to compare unilateral ALM inactivation trials to control trials (Fig. 5a), albeit without correcting for multiple comparisons (as ALM was the only region targeted for unilateral inactivation). When analysing effects of unilateral ALM inactivation, we combined left and right hemisphere laser ON trials. To address the imbalance in trial number between laser ON trials for each individual target site + epoch stimulation condition and the corresponding laser OFF control trials, we increased the observation weighting of the laser ON trials in the GLMM to achieve comparable trial balance across conditions.

To test whether unilateral inactivation resulted in spatial choice bias (Fig. 5b), we then restricted analyses to unilateral ALM laser ON trials, comparing the reduced model (Eq. 2) to a model that included a fixed effect term encoding whether the lottery location was ipsilateral to the photoinhibition-targeted hemisphere:

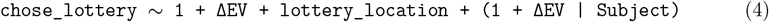

Because the output of the model was lottery choice, adding the lottery location term was equivalent to characterizing an ipsilateral choice bias. For this analysis we did not implement trial weighting as the relative numbers of trials in each condition were approximately balanced. For all GLMM analyses, cross-validation was repeated 50 times using different shuffled train-test splits, and reported *P* values are the medians across repetitions. GLMMs were fit using `fitglme` in Matlab using a binomial distribution with a logit link function and the Laplace approximation.

#### Mixed-agent model of optogenetic effects

To characterize the specific effects of optogenetic inactivations, we developed a mixed-agent model^21^ that used 5 parameters to transform the offers on each trial into a probability of choosing lottery as a weighted outcome of different types of agents (Fig. 3f): a rational agent, an abstract-value-bias agent, and a spatial-bias agent. Residual random choices unexplained by any agent are captured by a lapse parameter. We used this model, rather than the GLM-HMM model, for two main reasons. First, with the optogenetic experiment, the laser ON trials are expected to influence behaviour on that trial alone. This removes the core benefit of using an HMM: the assumption that mice switch from one state to another with low probability. The mixed-agent model treats trials as independent, so we can characterize the effects of the laser trials without concerns about state transitions. The second reason is that our mixed-agent framework allows us to estimate the full posteriors of a non-linear parametrization of behaviour. Risk-preference is typically modelled as utility curvature, which is a non-linear transform of value, so estimating this is outside the scope of GLM. Additionally, the effects of silencing on behaviour can be stimulus-dependent (i.e. horizontal shifts or changes in the slope of the psychometric curve), which are generally well captured by GLM, but they can also be stimulus-independent (i.e. vertical shifts or scaling of psychometric curves), which are not well captured by GLM.

The mixture model was fit using the following equation:

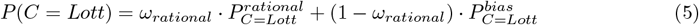

where *P* (*C* = *Lott*) is the probability of choosing the lottery as a mixture of the rational agents and bias agents, with *ω*_*rational*_ as the fraction of trials where animals are acting to maximize expected utility.

The rational agent is characterized by two parameters: *ρ, σ*^2^, where *U* = *V* ^*ρ*^: utility equals value (in normalized reward units) to the power of *ρ*. A concave utility function (*ρ* < 1) generates risk aversion, a linear function with *ρ* = 1 gives risk-neutral behaviour and a convex function (*ρ* > 1) generates risk-seeking behaviour^68^. We captured stochasticity in the rational agents choice by modelling the internal representation of expected utility, *EU* = *P · V* ^*ρ*^, as a Gaussian random variable.

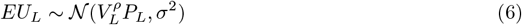

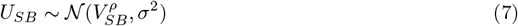

where the expected utility of lottery, *EU*_*L*_, and the utility of the surebet, *U*_*SB*_ are normal distributions. *V*_*L*_, *V*_*SB*_ refer to the magnitude of lottery and surebet reward and *P*_*L*_ is the probability of lottery payout. The probability of choosing lottery for the rational agent then becomes

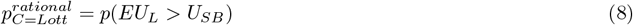

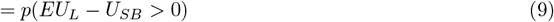

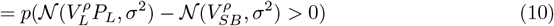

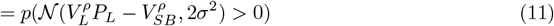

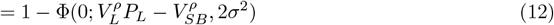

where 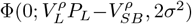 is the cumulative normal distribution with mean 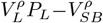, variance 2*σ*^2^ and evaluated at 0. Note that this provides fits with similar likelihood as the softmax choice function with *β* as temperature:

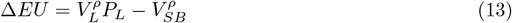

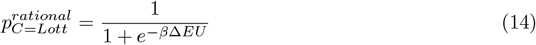

The contribution of the bias agents (and residual lapse) is characterizes as follows:

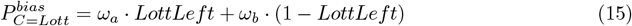

where

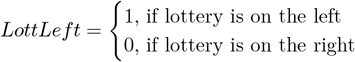

If ω_a_=ω_b_=.5,than 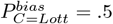 and we characterize this as lapse. If (*ω*_*a*_ = *ω*_*b*_)∧(*ω*_*a*_ ≫.5), then the bias is towards the lottery regardless of whether the lottery is on the left or right, so this would be characterized as a lottery bias. If (*ω*_*a*_ = *ω*_*b*_) ∧ (*ω*_*a*_ ≪.5), then the bias is towards the surebet regardless of whether the lottery is on the left or right, so this would be characterized as a surebet bias. If (*ω*_*a*_ = 1 − *ω*_*b*_) ∧ (*ω*_*a*_ ≫.5), then the bias is towards the left regardless of whether the lottery is on the left or right, so this would be characterized as a left bias. If (*ω*_*a*_ = 1 − *ω*_*b*_) ∧ (*ω*_*a*_ ≪ 5), then the bias is towards the right regardless of whether the lottery is on the left or right, so this would be characterized as a right bias. Cases where neither *ω*_*a*_ = *ω*_*b*_ nor *ω*_*a*_ = 1 − *ω*_*b*_ mean that there is a mixture of lapse with spatial and abstract biases.

In order to improve model convergence, we used 5 ‘raw’ parameters (*ϕ, ψ, ω*_1,2,3_) that were transformed into the variables described in the equations above as follows:

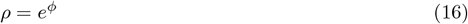

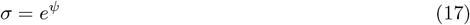

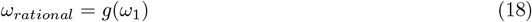

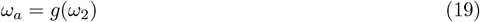

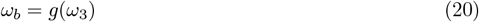

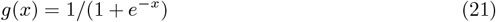

We fit *ϕ*, with prior 𝒩 (0, 0.5); *ψ*, with a prior of 𝒩 (− 3, 0.5); *ω*_1_, with a prior of 𝒩 (1, 1); *ω*_2,3_ with priors of 𝒩 (0, 1). The decision to have *ω*_1_ be shifted relative to the other *ω* was because mice were selected for sensitivity to Δ*EV*. We refer to these five parameters as *control* parameters, because they capture the behaviour on control (i.e., laser OFF) trials. We allowed each subject to deviate from each of these five population level parameters. For example, if we consider the utility curvature, *ρ* for mouse *i*,

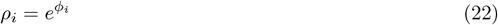

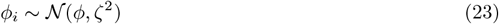

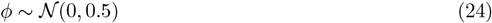

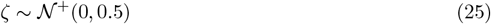

For all the subject level standard deviations we used a prior of ζ ~ 𝒩^+^(0, 0.5), a truncated normal distribution with lower bound=0.

We fit each inactivation dataset (unilateral and bilateral) separately. The trials used for the mixed-agent model were the exact same as the ones used for the GLMM analyses of effects of silencing ALM. For each dataset, we added two new parameters, reflecting the effects of first and second epoch silencing, for each raw parameter in order to estimate the effects of inactivation:

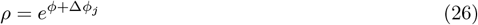

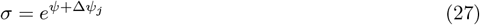

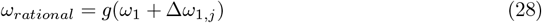

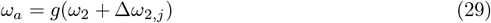

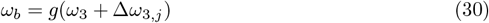

where Δ*ϕ*_*j*_ denotes the change in *ρ* in log space in the *j*th epoch (*j* = 1 for abstract choice, *j* = 2 for spatial planning), it had a prior of 𝒩 (0, 0.5); Δ*ψ*, with a prior of 𝒩 (0, 0.5), represents how the optoinhibition could shift noise; Δ*ω*_1,2,3_ (𝒩 (0, 1)) fit potential changes in *ω* before the logistic transformation, respectively, guaranteeing that *P* (*C* = *Lott*) remained a probability (i.e., bounded on [0,1]). We refer to these parameters as Δ parameters, because they capture the changes due to perturbation (Fig. 3g). Note, that all of the Δ parameters have priors with mean=0, so that our assumption is that the optogenetic silencing will have no effect on behaviour. This parametrization allowed us to treat the effects of perturbations as shifts of the raw parameters while guaranteeing that transformed parameters were constrained (e.g., *ρ* > 0, *σ* > 0, ∑*ω* = 1, ≤ 0 *ω*_*i*_ ≤ 1).

In order to simplify the reporting of the effects of perturbation in the results, we defined

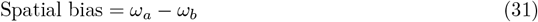

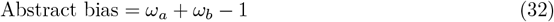

For unilateral inactivations, in order to combine left and right photoinhibition experiments into a single dataset, spatial bias was further transformed from left versus right biases into ipsi- or contralateral choice bias to the inactivated side (Fig. 5e,f).

We estimated the posterior distribution over model parameters with weakly informative priors using the brms (v2.22.0) R package^69^, a wrapper for Stan in R (`rstan`, v2.32.6)^70^. Stan is a probabilistic programming language that implements a Hamiltonian Monte Carlo (HMC) algorithm for Bayesian inference. Ten Markov chains with 24 000 samples each were obtained for each model parameter after 10 000 warm-up samples. The 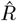 convergence diagnostic for each parameter was close to 1, indicating the chains mixed well.

#### Synthetic validation

To test the validity of our model, we created synthetic datasets with parameters that generated psychometric curves qualitatively similar to our perturbation data, which corresponded to changes in *ρ* or changes in *ω*. The mixed-agent model was fit to the synthetic datasets, and was able to recover the generative parameters accurately (Fig. S8a).

### Electrophysiology

#### Surgery for acute recordings

Post learning, mice were habituated to the recording setup for 1–3 weeks, until behavioural performance recovered in the recording setup. Mice then underwent a second, minor surgery to prepare for acute electrophysiological recordings. Mice were anaesthetized with isoflurane (1.5%) and placed in a stereotaxic apparatus (Kopf Instruments). The surface of the Super-bond used for headplate implantation was cleaned with Hibiscrub followed by sterile saline. Two craniotomies (~ 2 mm diameter) were made bilaterally over ALM (AP +2.5 mm, ML ± 1.5 mm from bregma). An additional cranial opening (~ 0.5 mm diameter) was drilled above the cerebellum to secure the ground wire to the skull. Each craniotomy was covered with Dura-Gel (Cambridge NeuroTech), and a plastic protective cap was placed over the headplate to shield the exposed areas and maintain sterility between recording sessions.

#### Recording

For electrophysiology recordings in awake behaving animals, neural activity was recorded using Neuropixels 1.0 probes (IMEC, Belgium), digitized at 30 kHz, acquired through a PXIe-1083 chassis (National Instruments), and saved using SpikeGLX (https://github.com/billkarsh/SpikeGLX).

For electrophysiological validation of photoinhibition in the laser-scanning experiments, we used a 64-channel silicon probe (ASSY-236-H9, Cambridge NeuroTech) to minimize photoelectric artifacts. Neural signals were digitized at 30 kHz, amplified and band-pass filtered (0.6 Hz to 7500 Hz) using a 64-channel Intan headstage, and processed, visualized and saved using the Open Ephys acquisition board (https://open-ephys.github.io/acq-board-docs/) and GUI.

Before each recording session, the plastic headplate cap was gently removed to expose the Dura-Gel sealed craniotomies. A microscope (KAPS) was used to inspect the craniotomy and guide probe insertion. Probes were lowered at 7 µm*/*s using programmed automatic micromanipulators (Sensapex), followed by a 10 minute settling period to minimize neural signal drift during recording.

During recording, at the end of each trial, a serial TTL message encoding the current trial number was sent from behavioural control hardware to the acquisition system to synchronize the neural signal with the behavioural data. At the end of each recording session, probes were retracted at 15 µm*/*s, the craniotomies were re-covered with a sterilized plastic cap, and the animal was returned to its home cage. Each behavioural animal was recorded for 3 to 7 sessions, depending on performance. Animals used for electrophysiological validation of photoinhibition (see below) were recorded for 1 to 2 sessions.

#### Electrophysiological data preprocessing

For electrophysiological signals recorded using Neuropixels 1.0 probes, raw data were pre-processed with CatGT (https://billkarsh.github.io/SpikeGLX/help/catgt_tshift/catgt_tshift/). A *tshift* correction was applied, and AP-band data were filtered using a 12th-order Butterworth band-pass filter (300 Hz to 9000 Hz). A global common-average reference (CAR) was applied to reduce artifacts shared across channels.

Offline spike sorting for both CatGT-processed Neuropixels data and 64-channel silicon probe recordings was performed using Kilosort v4^71^ for each recording session. Manual curation was conducted in Phy 2.0 (https://github.com/cortex-lab/phy). Units were classified as putative neurons only if the waveforms exhibited a clearly spatial decay across adjacent probe channels. Units that showed low percentage of missing spikes, low refractory period violations, and minimal drift across time were marked as single-units, and other units were marked as multi-units. Units showing substantial drift that resulted in a clear loss of spikes over time were excluded.

Quality metrics and spike waveform features were computed using SpikeInterface^72^. We included both single-units and multi-units from manual curation if the presence ratio (proportion of discrete 1 min time bins in which at least one spike occurred) exceeded 0.95 for the entire session and the mean firing rate exceeded 0.5 Hz. To ensure stability across the recording, we imposed three additional criteria, inspired by similar metrics in previous work^73^: the rolling 15 min firing rate (computed with a 2 s step) could not fall below 40% of the mean firing rate across the session; the rolling 10 min firing rate could not fall below 50%; and the rolling 5 min firing rate could not fall below 50%. Units that met all of the above criteria were included in subsequent analyses.

### Electrophysiology analyses

#### Single-neuron analyses

For the example neuron raster plots and peri-stimulus time histograms (PSTHs; Fig. 4a), spike times were aligned to the onset of the lottery sound cue and analyzed over a 3 s window (from 0.5 s before the sound cue onset until 0.5 s after the lick spout extension). The spike trains were binned at 10 ms resolution and smoothed with a causal half-Gaussian kernel (*σ* = 100 ms).

To characterize the temporal dynamics of task-related variable encoding in these example neurons (Fig. 4b), we fit a linear regression model to continuous time bins of z-scored firing rate using MATLAB’s `fitlm`, as the following:

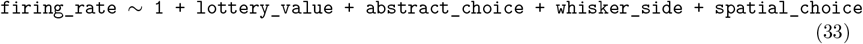

where `lottery_value`is a continuous variable representing the five lottery values, `abstract_choice` is a binary indicator of the animal’s abstract choice (1 = lottery chosen, 0 = surebet chosen), `whisker_side` is a binary indicator of the whisker stimulation side (1 = whisker stimulation delivered contralateral to the recording hemisphere, 0 = ipsilateral), and `spatial_choice` is a binary indicator of the animal’s upcoming directional licking response (1 = lick contralateral to the recording hemisphere, 0 = ipsilateral).

To quantify single-neuron task representation across the population over the entire first or second epoch (Fig. 4c,d), we applied the same linear model to neural activity extracted from the lottery value epoch (0 s to 1 s after lottery cue onset) or motor planning epoch (1 s to 2 s after the lottery cue onset). We used a threshold of *P* < 0.05 for the coefficients to identify significantly task-selective neurons, and *χ*^2^ tests to determine whether the proportions of distinct types of selective neurons were statistically different. Similar analyses were used to characterize layer-specific ALM sub-populations (Fig. S9d,e).

#### Population decoding

To quantify task-relevant information on the population level, we performed cross-validated decoding analyses using ALM pseudopopulations over continuous time bins (Fig. 4e-h). Spike trains were converted into PSTHs using a causal smoothing kernel (square window, 200 ms width, 10 ms step). PSTHs were aligned to the lottery sound onset and analyzed over a 3 s window (from 0.5 s before the sound cue onset until 0.5 s after the lick spout extension), yielding 300 time bins (*T*). To test how population size influenced decoding, we generated pseudosessions (200 for each population size) with either 100, 500, or 1000 neurons by sampling neurons without replacement from the full dataset. Neurons were excluded if they did not satisfy the balancing criteria described below.

We decoded four task variables: lottery value, abstract choice, whisker side and spatial choice, defined similarly to single-neuron regression analyses. The binary variables jointly define the 4 choice trial types. For lottery value decoding, we subsampled trials so that there were equal numbers of abstract choice, whisker side, and spatial choice trials within each lottery value trial type. This guaranteed that lottery value information was not confounded by any other binary variable, such as abstract choice. The resulting pseudosession dataset formed a tensor size of 20 × *N* × *T* (trials × neurons × time bins). For all the other binary variables decoding, we subsampled 6 trials per trial type for the 4 trial types (24 total). We also balanced the number of lottery value trials within each of the 4 trial types to remove lottery value information when decoding the other 3 binary variables. This generated tensor size of 24 × *N* × *T* for decoding.

To remove noise correlations between simultaneously recorded neurons for pseudopopulation decoding, firing rates of each neuron were z-scored across trials independently at each time bin. We then conducted cross-validated decoding (4 folds for lottery value decoding and 6 folds for binary variable decoding). For each fold, principal component analysis (PCA) was used on z-scored training data for dimensionality reduction and de-noising, and the top 5 PCs were included. Both training and test data were then projected onto this low-dimensional subspace to generate *X*_*train*_ and *X*_*test*_. We used linear regression (Matlab function `glmfit`) to estimate the 5 × 1 matrix *B* as the best direction to separate the different lottery values as follows:

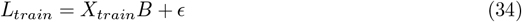

where *L*_*train*_ is the true lottery value for the training set. Then we estimated the lottery values on the left out trials

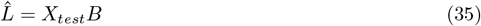

The decoding performance was quantified using the Pearson’s correlation between the true lottery values and the cross-validated predicted values 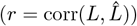 for all trials. All the other binary variables were decoded using a linear SVM classifier (MATLAB function fitcsvm with a linear kernel) as follows:

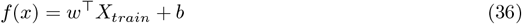

with test labels predicted using

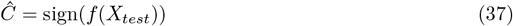

and the decoding accuracy was computed as the fraction of correctly classified trials.

For each pseudosession and time bin, we generated a null distribution by shuffling trial labels 1000 times and computing the resulting chance decoding accuracy. If the actual decoding accuracy in a particular time bin exceeded the 99th percentile of the shuffled distribution, activity in that time bin was considered to contain significantly above-chance task information. Across all the pseudosessions, a time bin was considered significant if ≥ 50% of pseudosessions exceeded this threshold. All analyses were conducted using MATLAB R2022a (Mathworks) on an high-performance computing cluster.

#### Noise correlation analyses

To characterize the functional connectivity between neural populations with specific tuning properties within and across hemispheres, we calculated trial-by-trial noise correlations between pairs of simultaneously neurons during the first or second task epoch (Fig. 6f,g). We removed signal correlation by subtracting mean activity of each trial type from the per trial spike count for every neuron before calculating across-trial correlations between neuron pairs. In other words, noise was calculated as the trial-by-trial deviation from the mean activity for that trial type. Noise correlations were computed only for pairs of significantly task-selective neurons recorded simultaneously (see single-neuron analyses for methods of identifying task-selective neurons).

For noise correlation analyses during the first task epoch, we included all simultaneously recorded neurons with significant abstract choice encoding, classified as lottery- or surebet-preferring based on whether their firing rates were higher for lottery or surebet choices. For the second task epoch, we only included pure-selectivity action neurons that had significant spatial action encoding but not any other task-relevant information. Within this group, neurons were classified as ipsilateral- or contralateral-preferring based on whether their firing rates were higher for ipsilateral or contralateral upcoming actions relative to the recording side. All simultaneous bilateral ALM recording sessions (n = 2 mice, 3 sessions per mouse) were included in these analyses. The noise correlation values were pooled across sessions.

### Electrophysiological validation of photoinhibition

To characterise the spatiotemporal resolution of optogenetic perturbations on neural responses, we performed acute electrophysiological recordings during laser stimulation in two awake VGAT-ChR2-EYFP mice. A 1.5 mm craniotomy was made over ALM (centred at AP 2.5, ML 1.5 mm from Bregma) for the insertion of a single-shank 64-channel silicon probe (ASSY-236-H9, Cambridge NeuroTech). Across sessions, the probe was lowered to different depths, enabling recordings from 200 − 2400 µm from brain surface, so that we can sample neurons across cortical layers. In one VGAT-ChR2-EYFP mouse we also performed recordings from visual cortex (craniotomy centred at AP −2.9, ML 1.4 mm from Bregma) to confirm optogenetic effects were similar across cortical regions (Fig. S6c).

Optogenetic stimulation was delivered using the Zapit laser system^36^ as described above (473 nm laser, OBIS LX, Coherent). The focal point was calibrated before each session, and we stimulated 4 locations along the anterior–posterior axis covering a 0 − 3 mm range at 1 mm intervals from the recording site (Fig. S6a). Laser stimulation was 40-Hz sinusoidal waveform lasting for 1 s, with a 200 ms linear rampdown at the end of the stimulation window to minimise rebound excitation. Laser ON and laser OFF (control) trials were randomly interleaved (~ 400 trials/session). We also tested multiple laser powers (1, 1.5, 2, and 4 mW) in an interleaved design. We combined 1.5 and 2 mW conditions for analyses since they evoked similar effects (Fig. S6d,e).

Spike sorting was performed using Kilosort v4^71^, followed by manual curation in Phy 2, as described above. Putative excitatory units were identified as neurons that did not increase firing rate with 4 mW laser stimulation. Fig. S6b shows the spike rasters and PSTHs for one example putative excitatory unit for 2 mW and 4 mW optogenetic stimulations at different stimulation locations. Spike times were aligned to laser onset, spanning a 3 s window (1 s before laser onset to 1 s after laser offset). To generate the PSTHs, spike times were counted over 10 ms time bins and smoothed using a causal half-Gaussian kernel (*σ* = 150 ms).

To characterize effects of optogenetic silencing on the population level, for each unit and stimulation condition, firing rates were averaged within the 1 s laser stimulation period and normalized relative to pre-laser baseline activity (1 s). Putative excitatory units were then pooled to quantify the average effects for each stimulation condition (Fig. S6c,d). Rebound activity was calculated as percent firing rate change in the 500 ms window following laser offset, relative to the pre-laser baseline activity (500 ms). Kernel density and median value of rebound effects across the population were plotted for each stimulation condition (Fig. S6e).

### Dynamical models

We developed a dynamical rate model of ALM to test how interhemispheric interactions can facilitate the transformation from abstract economic choices to spatial actions, and give rise to the distinct effects of silencing ALM across epochs (Fig. 6). We used 6 populations made of 30 neurons each (3 populations per hemisphere): each hemisphere had three distinct types of population, ‘lottery’ neurons,ℒ ‘surebet’ neurons, 𝒮 and motor neurons, ℳ We denote the hemisphere of a population using a subscript *L, R*. Each neuron i in population *α* follows the update rule:

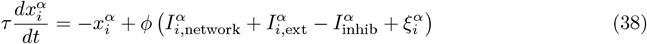

with *dt* = 0.02s, *τ* = 0.16*s ϕ*(*x*) = *max*(0, *tanh*(*x*)), and 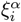 is Gaussian noise ~ 𝒩 (0, 0.11).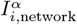 is the input from the other neurons in the network:

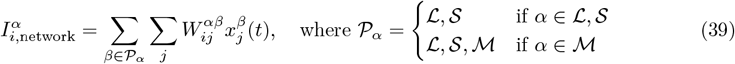

the weights are sampled from the normal distribution 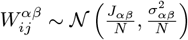 where 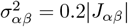 controls the heterogeneity, and *J*_*αβ*_ represents the mean connection strength between population *β* and population *α*. The structure of *J*_*αβ*_ is defined by the functional role of the connected populations, as shown in Tab. 2.

**Tab 2:**
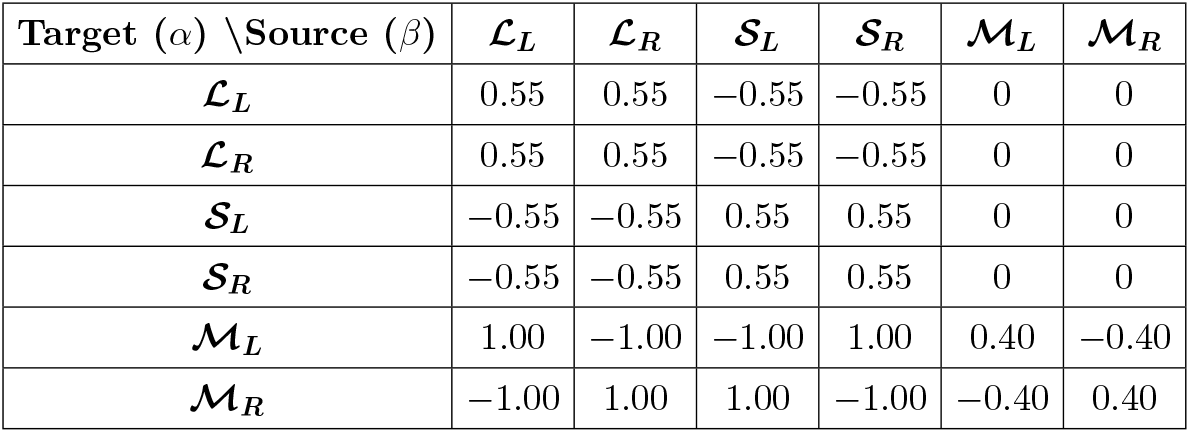
Weight matrix.

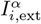 is the external input

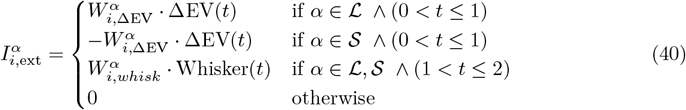

Where 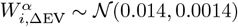, 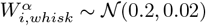 The strength of optogenetic inhibition,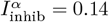 The inhibition was applied bilaterally and unilaterally in the first and second epoch.

The final action was taken by following a softmax policy, where the probability of choosing to lick right was defined as:

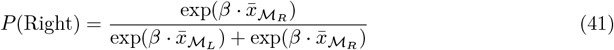

where 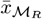 and 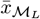 are the mean activity in the 200ms interval after the end of the second epoch of the right and left motor populations, and *β* = 20 is a temperature parameter, controlling the steepness of the psychometric curve.

The psychometric curves were generated using 17 ΔEV values ranging from −1 to 1, with 200 trials per condition. The noise correlations were computed using 5 ΔEV values and 20 trials per condition. Simulation was done using Python with the numpy package^74^.

## Author contributions

O.M.G. and C.A.D. conceived the project, with J.C.E.’s input. O.M.G. and C.A.D. designed the behavioural setup and task. J.L. contributed to the electronic design of the setup. O.M.G., C.B., and J.T. performed surgeries. O.M.G., C.B., J.T., Y.P., and N.Z. contributed to behavioural training. J.L., J.T., and N.Z. performed histology. O.M.G. collected and analysed widefield and optogenetic data, with J.T. and Y.P.’s assistance. C.B. and J.L. built the acute Neuropixels recording setup. C.B. collected and analyzed the electrophysiological data. C.B. and J.C.E. implemented the mixed-agent models. T.P.H.S. conducted the decoding analyses for widefield experiments. J.W. implemented the early versions of GLM-HMM. G.B. and J.L. performed the dynamical modelling, with C.C. and J.C.E.’s inputs. C.A.D. and J.C.E. provided supervision and inputs to all data analyses. C.A.D. and O.M.G. wrote the manuscript, with contributions from co-authors.

## Acknowledgments

This work was funded by the Sainsbury Wellcome Centre Core Grant from the Gatsby Charitable Foundation (GAT3755) and Wellcome (219627/Z/19/Z); and by a Wellcome Early Career Grant (309104/Z/24/Z) awarded to O.M.G. We thank members of the Duan laboratory and members of the Erlich laboratory for insightful discussions and advice; P. Zhang for assisting with acute recording experiments; T. Margrie, T. Mrsic-Flogel, K. Harris, T. Behrens, P. Coen, and T. Branco for comments on the manuscript; SWC Neurobiological Research Facility and M. Li for animal support; and R. Barrett, D. Halpin, S. Townsend, and S. Hammett from FabLabs for contributions to hardware design and setup.

## Code and Data Sharing

Code and data related to this manuscript will be available at the time of publication at https://gitlab.com/sainsbury-wellcome-centre/delab/publications/alm-riskychoice-2026. To get early access, please contact the corresponding author.

---

**Fig. S1:**
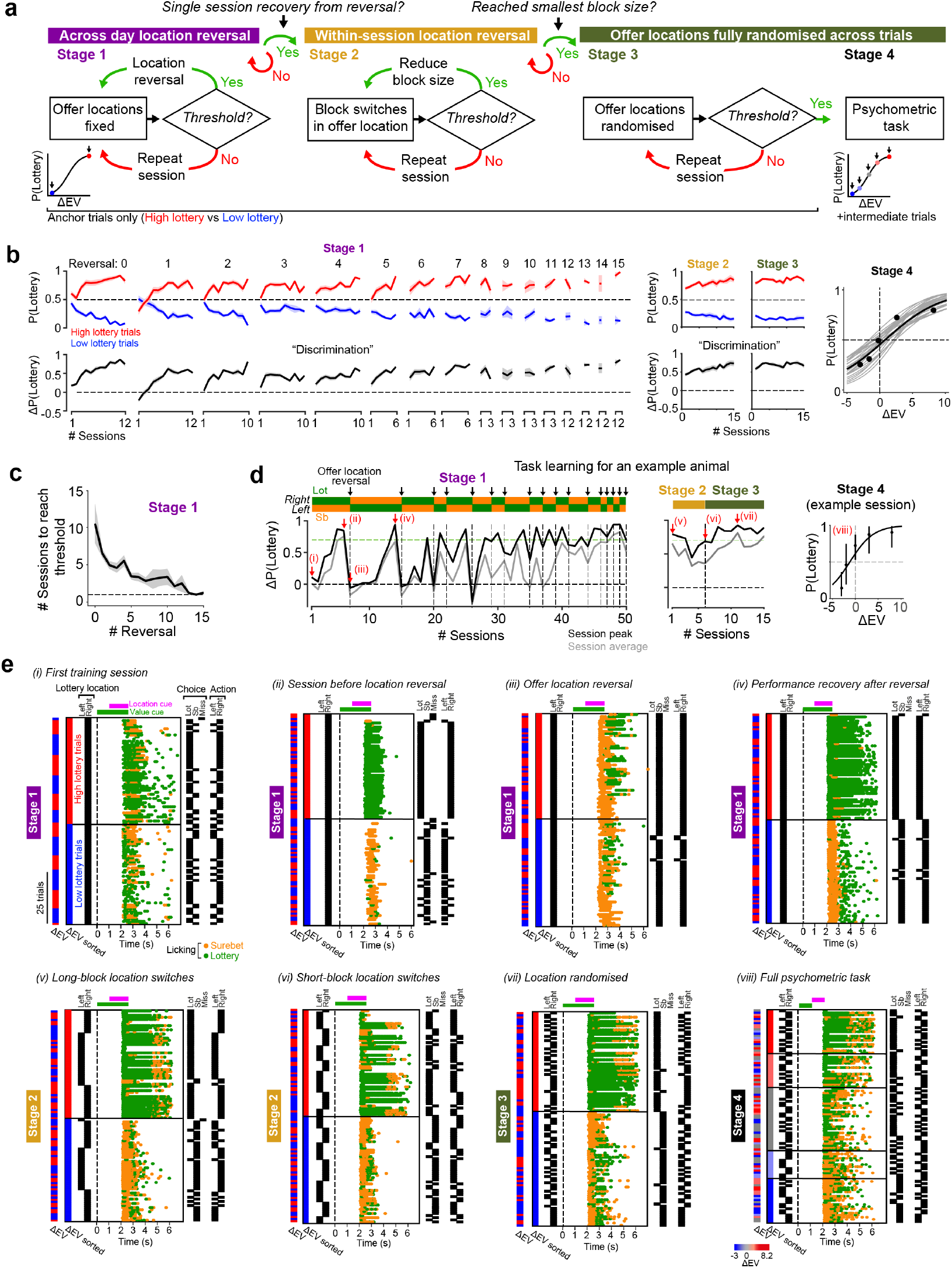
Behavioural training and task learning. **a**, Overview of training curriculum. **b**, Performance across learning stages. Stage 1: across-session reversals in offer location. Top, P(Lottery) for high (red) and low (blue) ΔEV trials. Bottom, discrimination across trial types, ΔP(Lottery). Data show the mean and s.e.m. across mice (n = 27). Stage 2, within-session block-switches in offer location. Stage 3, offer locations randomised within-session. Stage 4, Full task performance. Grey, psychometric curves for individual mice; black, concatenated data across mice. Data show session peak performance as assessed across a 100-trial rolling window. **c**, Number of sessions to recover performance following repeated offer-location reversals in Stage 1. Thick line and shading, mean ± s.e.m across mice (n = 27). **d**, Learning trajectory for an example animal. For Stages 1-3, session peak (black) and session average (grey) ΔP(Lottery) is shown across training days. For the example Stage 4 psychometric session, data points show mean and 95% binomial CI. **e**, Lick raster plots for the example sessions highlighted with red arrows in **d**. For each plot, trials are sorted by ΔEV along the y-axis, with the true ΔEV trial sequence indicated on the far left. The lottery location is indicated by black ticks on the left of *t* = 0 s. Abstract choice and spatial choice for each trial are indicated on the far right of each plot. Individual licks to the lottery and surebet lick spout are colour coded green and orange respectively. The green and magenta bars at the top of the axes indicate the timing and duration of lottery value (sound) and location (whisker) cues respectively. Each plot shows 100 consecutive trials identified as the session peak performance.

**Fig. S2:**
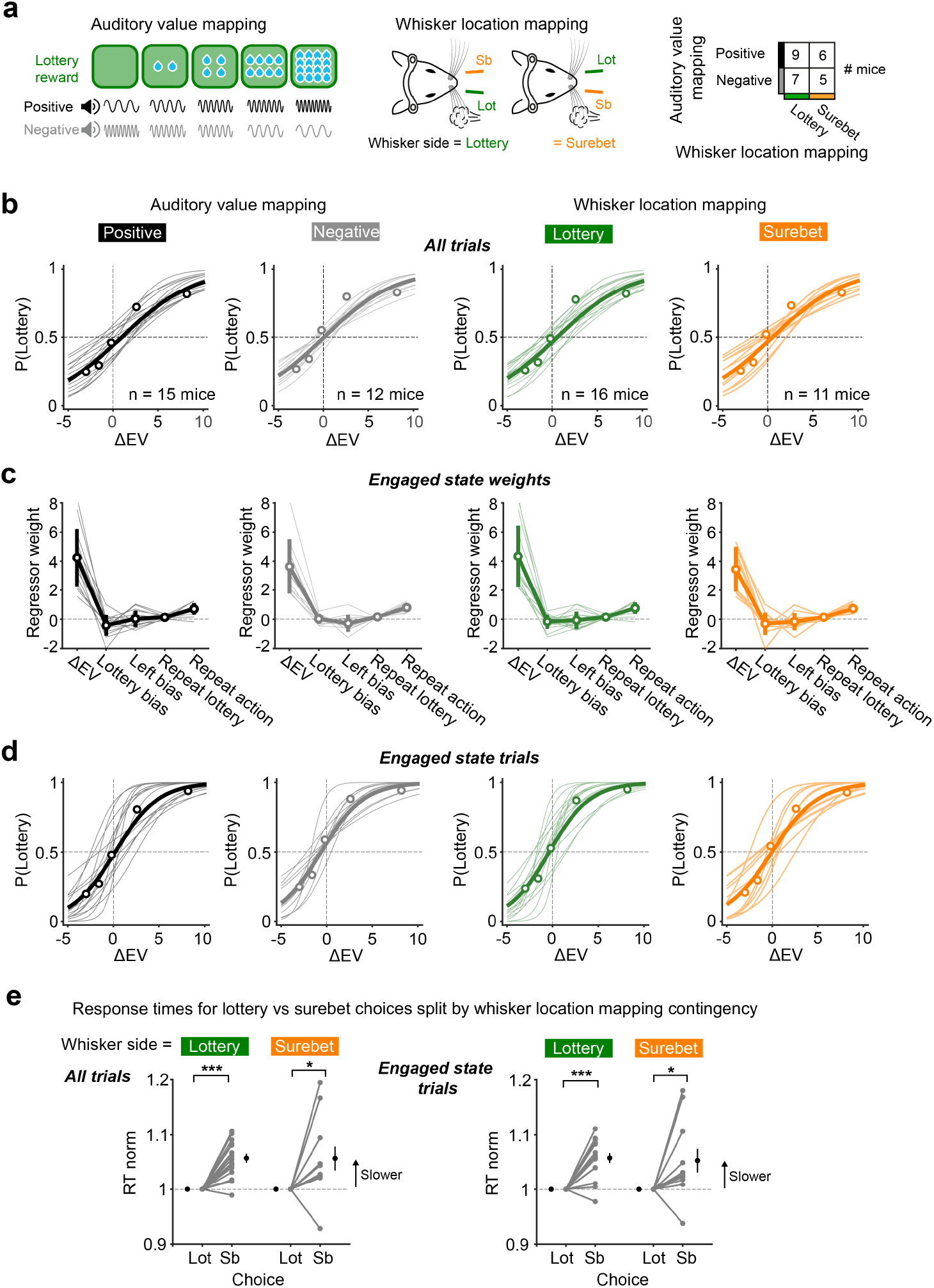
Lottery value and location stimulus contingencies. **a**, Left, mice were either trained with a positive auditory-to-value mapping (e.g. high pure tone frequency signals large lottery reward) or a negative mapping (e.g. high tone frequency signals low lottery reward). Middle, the side of whisker stimulation corresponded to the lottery offer location for some mice, and the surebet offer location for others. Right, numbers of trained mice in each stimulus contingency group. **b**, Task performance across stimulus contingencies. Thin lines show logistic fits for individual mice, thick lines and data points show average fits and data concatenated across mice. **c**. Inferred GLM weights during the engaged state (from 3-state GLM-HMM). Data points indicate mean ± s.e.m.; individual lines represent individual mice. **d**, Behavioural performance on engaged trials, compared to **b** for all trials. **e**, Response times (RTs) for lottery and surebet choices across mice with different whisker location mappings. Data show median RTs normalised to lottery choices. Grey, individual mice, black, mean ± s.e.m. Paired comparisons were Wilcoxon signed-rank tests. Significance is indicated as \**P* < 0.05, \*\**P* < 0.01, \*\*\**P* < 0.001.

**Fig. S3:**
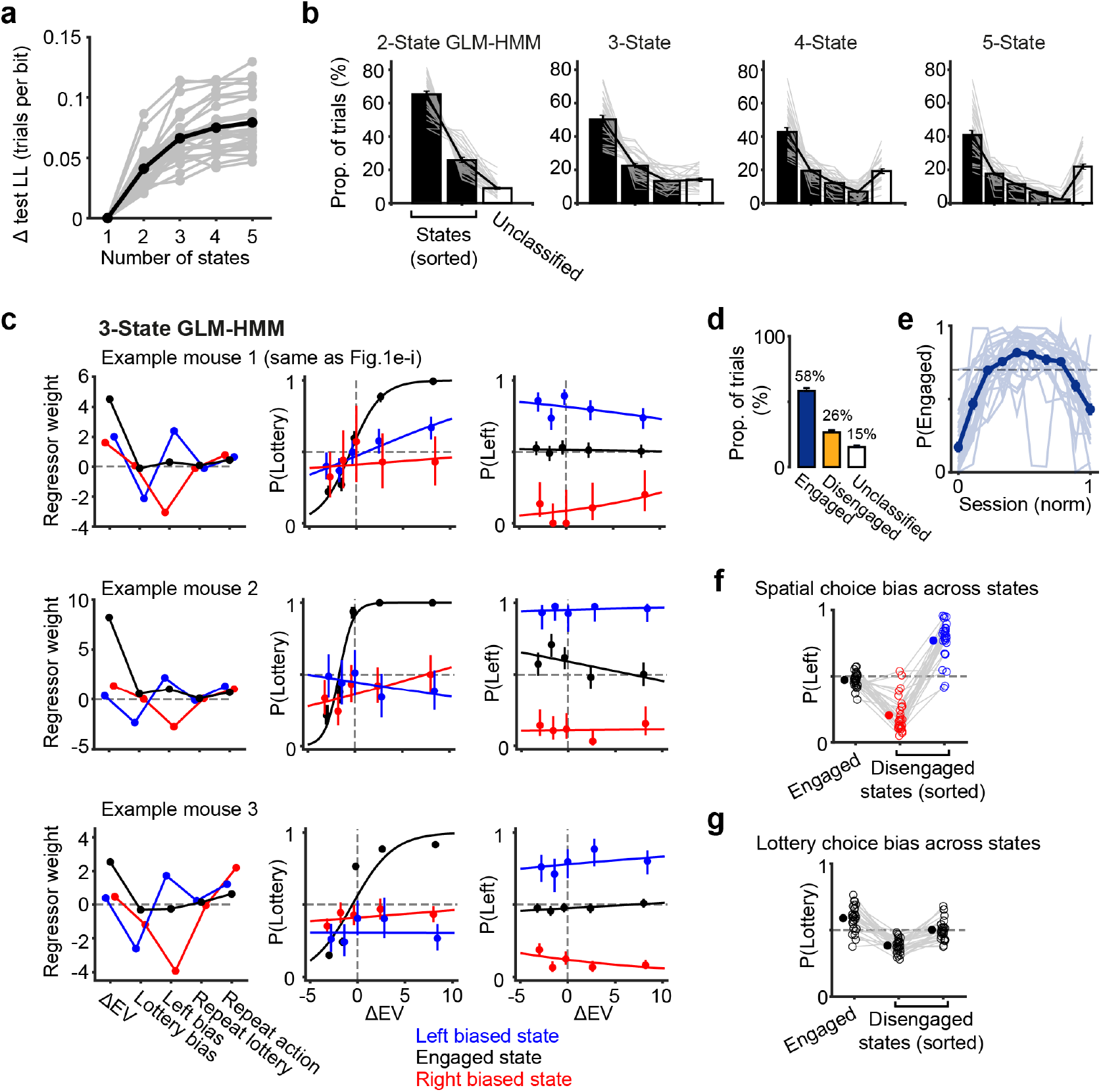
A GLM-HMM captures engaged and spatially biased behavioural states. **a**, Model comparison across GLM-HMMs with varying numbers of latent states. Change in per test trial log-likelihood relative to a 1-state GLM is shown. The black line shows the mean across mice (n = 27), grey lines show individual mice. Model performance was assessed on held-out data (5-fold cross-validation across sessions). **b**, Proportion of trials assigned to each state across models. Trials were classified using a P(State) threshold of 0.7; unclassified trials (P(State) < 0.7 for all states) are shown in white. Bars indicate the mean across mice, error bars show s.e.m (n = 27 mice). Thin grey lines indicate individual mice. State data from each animal were first sorted in descending order. **c**, Example model and behaviour data from three mice for the 3-state GLM-HMM. Left: inferred GLM regressor weights. Middle, lottery choice behaviour across states. Right, leftward choice behaviour across states. For each mouse, the model identified a task-engaged state with a high ΔEV weight (black) and two ‘disengaged’ states showing strong leftward (blue) and rightward (red) spatial choice biases respectively. Data points show the mean ± 95% binomial CI. **d**, Average proportion of trials assigned to engaged and disengaged states. Data shows mean *±* s.e.m. **e**, Probability of being in the engaged state across trials in a session. Individual lines show individual mice. **f**, Mean spatial choice bias across all trials in each state in the 3-state GLM-HMM. During the engaged state, mice showed minimal spatial choice bias. However, choices during the two disengaged states were characterised by opposing directional biases. Open markers and thin grey lines indicate individual mice. Filled markers show the mean across mice (n = 27). **g**, Similar to **f** but showing lottery choice bias across states. Strong biases in lottery choice were not a consistent feature of disengaged state performance.

**Fig. S4:**
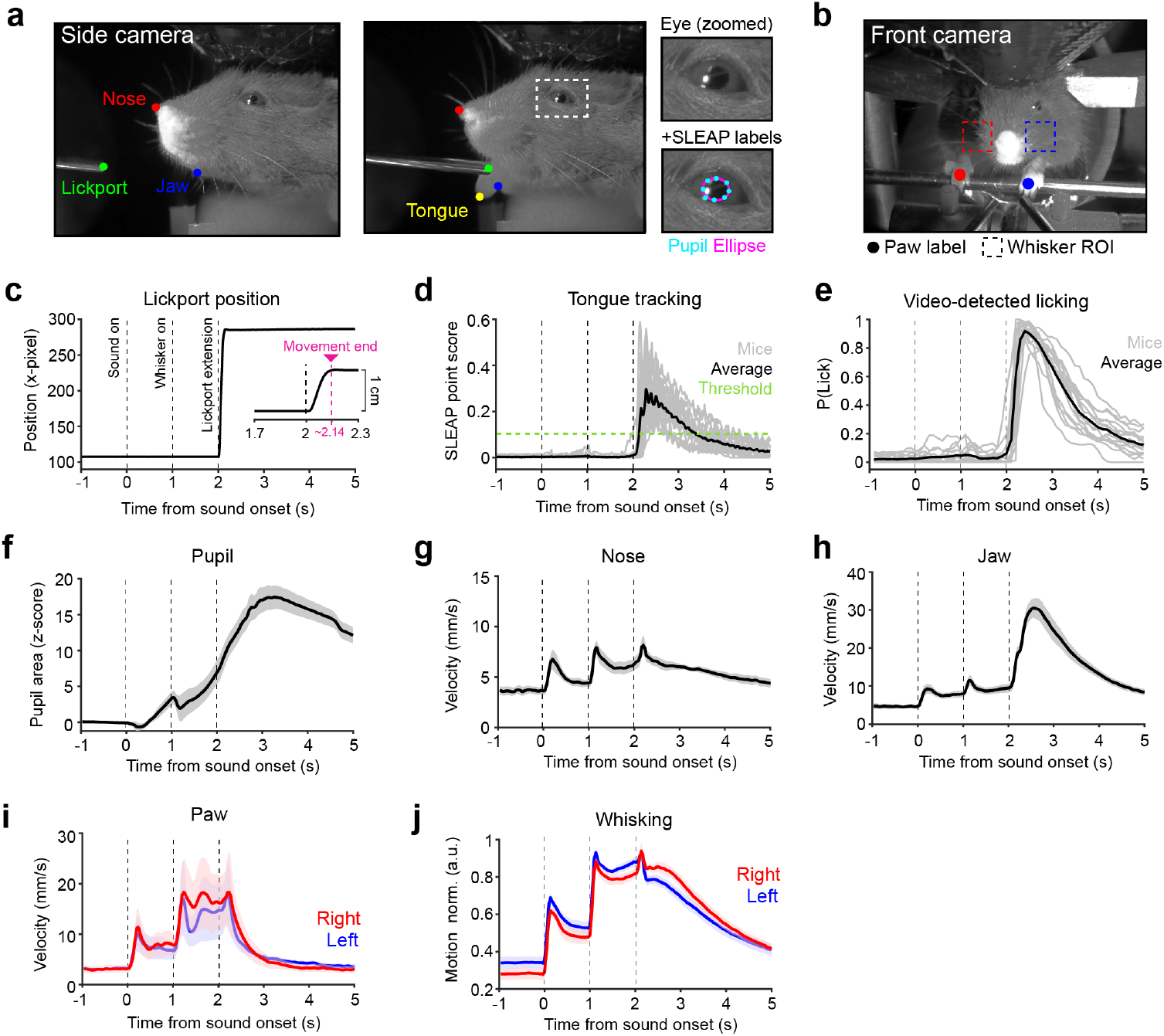
Behavioural videography. **a**, Example images from the side camera showing the lickport retracted (left) and extended (middle). A close up of the eye (white dashed box) is shown on the right. SLEAP nodes are shown as coloured circular markers, note that the tongue node (yellow) is only present during licking (middle image). For pupillometry, an ellipse (magenta) was fit to 8 SLEAP nodes outlining the pupil. **b**, Example image from the front camera. SLEAP nodes for the left (blue) and right (red) forepaws are shown as circular markers. Regions of interest (ROI) around the left (blue) and right (red) whisker pads are shown as dashed boxes. **c**, Timing of lickport extension quantified as the lateral position of ‘lickport’ node across time. The inset shows a close-up of the translation period indicating the time to reach final position is short (~140 ms). Data show the mean for an example session. **d**, Videography detection of the tongue using the side camera. The point-score for the tongue node provides a measure of SLEAP tracking confidence of tongue detection across time. Thin lines show individual mice (n = 16 mice; 1 representative session per mouse), black line shows the average across mice. The green line shows the point-score value used as a binary detection threshold. **e**, Quantification of licking across the trial. Using the point-score threshold in **d**, lick probability was assessed in 100 ms bins across each trial. Thin lines show individual mice (n = 16), black shows the average. Mice showed minimal anticipatory licking outside of the response window. **f**, Average pupillometry response across the trial. Data show mean ± s.e.m. across 6 widefield mice. **g**, Average nose movement across the trial. Data show mean ± s.e.m. across mice. **h**, Similar to **g**, but showing jaw movement. **i**, Similar to **g**, but showing left (blue) and right (red) paw movement. **j**, Average left (blue) and right (red) whisker motion. Data show mean ± s.e.m. across mice.

**Fig. S5:**
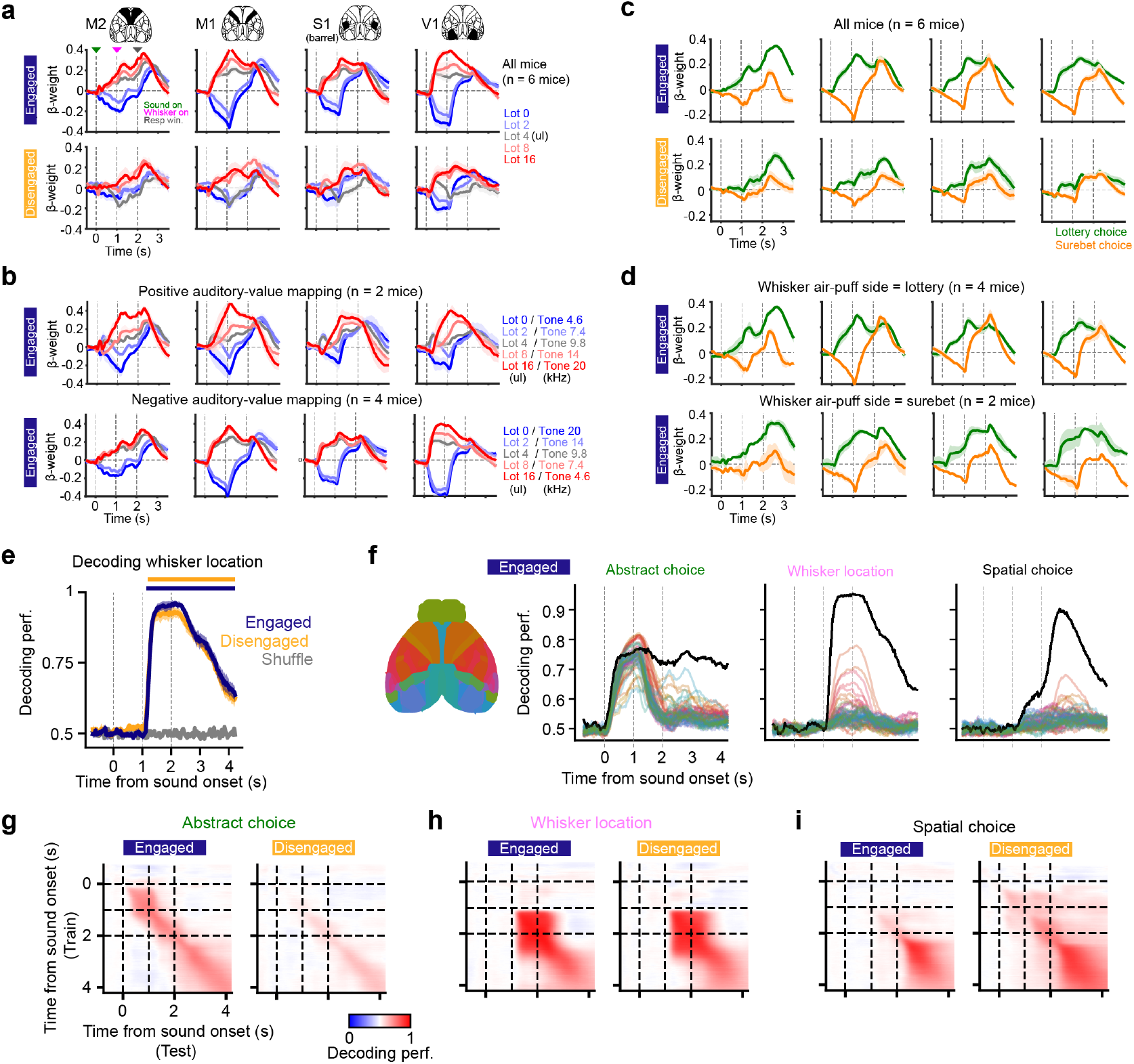
Comparison of widefield signals across behavioural states and stimulus contingencies. **a**, Lottery value encoding across behavioural states. Kernel regressor traces were averaged across different bilateral cortical areas. Top, data from engaged trials; bottom, data from disengaged trials. Data show mean ± s.e.m across mice. **b**, Lottery value encoding in mice trained with opposing auditory-to-value mappings. Kernel regressor traces were averaged across cortical areas as in **a**. Top, data from positive tone-frequency-to-value mapping mice (n = 2); bottom, data from negative tone-frequency-to-value mapping mice (n = 4). Data show mean ± s.e.m across mice, engaged trials only. **c**, Abstract choice encoding across behavioural states. Kernel regressor traces for lottery (green) and surebet choice (orange) across cortical areas as in **a**. Top, data from engaged trials; bottom, data from disengaged trials. Data show mean ± s.e.m across mice. **d**, Abstract choice encoding across mice trained wtih opposing whisker-to-offer location rule mappings. Kernel regressor traces for lottery (green) and surebet choice (orange) were averaged across cortical areas as in **a**. Top, data from mice where whisker stimulation side cued lottery offer location (n = 4 mice). Bottom, data from mice where whisker stimulation side cued the surebet offer location (n = 2 mice). Data show mean ± s.e.m across mice, engaged trials only. **e**, Decoding of whisker stimulation side using fluorescence activity across all CCF-defined cortical areas across engaged and disengaged trials. Data show mean and s.e.m across mice (n = 6). Bars on top indicate time points when decoding is significantly better than chance tested using Wilcoxon signed-rank tests and *P* < 0.05. Decoding performance for whisker location was not significantly different between the engaged and the disengaged states. **f**, Decoding of task variables using single CCF-defined cortical regions (thin coloured lines) vs all cortical regions (thick black line) in engaged trials. **g**, Cross-temporal and cross-validated decoding of abstract choice using multi-region widefield fluorescence signals in engaged and disengaged trials. Data show mean across 6 mice. **h**, Similar to **g**, for whisker stimulation side. **i**, Similar to **g**, for spatial choice.

**Fig. S6:**
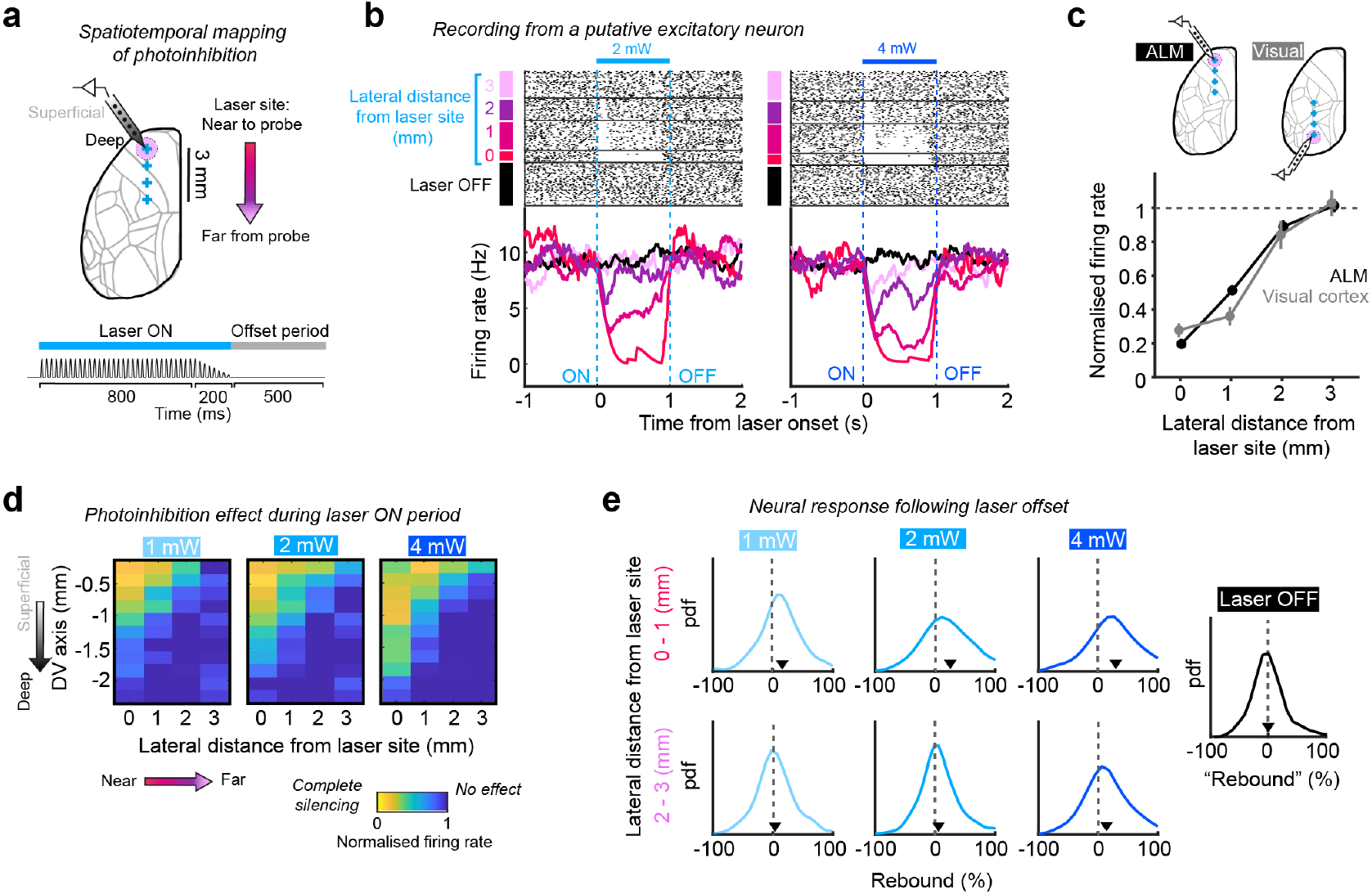
Electrophysiological validation of optogenetic silencing effects. **a**, Schematic for electrophysioloigical validation experiments in VGAT-ChR2-EYFP mice. Top, neural activity was recorded in awake mice using Cambridge Neurotech silicon probes. Spread of optogenetic silencing effects was characterized along the DV axis for different cortical depths (across the probe shank), and multiple lateral distances (targeting the laser at 0, 1, 2, or 3 mm away from the recording site). Bottom, laser profile used in acute recording experiments, similar to what was used in behavioural experiments. **b**, Example spike raster (top) and peristimulus time histogram (PSTH, bottom) of a putative excitatory neuron recorded in ALM, on laser OFF or laser ON trials at 2 mW (left) and 4 mW (right) laser powers. Laser ON trials were sub-grouped according to different lateral distances between the laser beam and the recording site. The example neuron showed distance- and laser-power-modulated silencing of cortical activity, with minimal rebound after laser offset. **c**, Average suppression of activity across lateral distances between the laser beam and the recording site from optogenetic validation experiments performed in ALM (black) and visual cortex (grey). Data show mean and s.e.m. across n = 117 putative excitatory cells recorded in ALM and n = 23 putative excitatory cells recorded in visual cortex across 2 mW and 4 mW laser trials. Data are from the same VGAT-ChR2-EYFP mouse. **d**, Average suppression of activity across cortical depths and lateral distances between the laser beam and the recording site, for 1 mW (left), 2 mW (middle), and 4 mW (right) laser powers. Data show mean across n = 354 putative excitatory cells recorded in ALM (n = 2 VGAT-ChR2-EYFP mice). **e**, Distribution of post-laser offset rebound in firing rate of putative excitatory neurons in ALM for different laser powers and lateral distances. Rebound was calculated as percentage of firing rate change during the 500 ms following laser offset, relative to pre-laser baseline period. This metric was also calculated using the same analysis windows for laser OFF trials for comparison (black). Arrows indicate the median of each distribution.

**Fig. S7:**
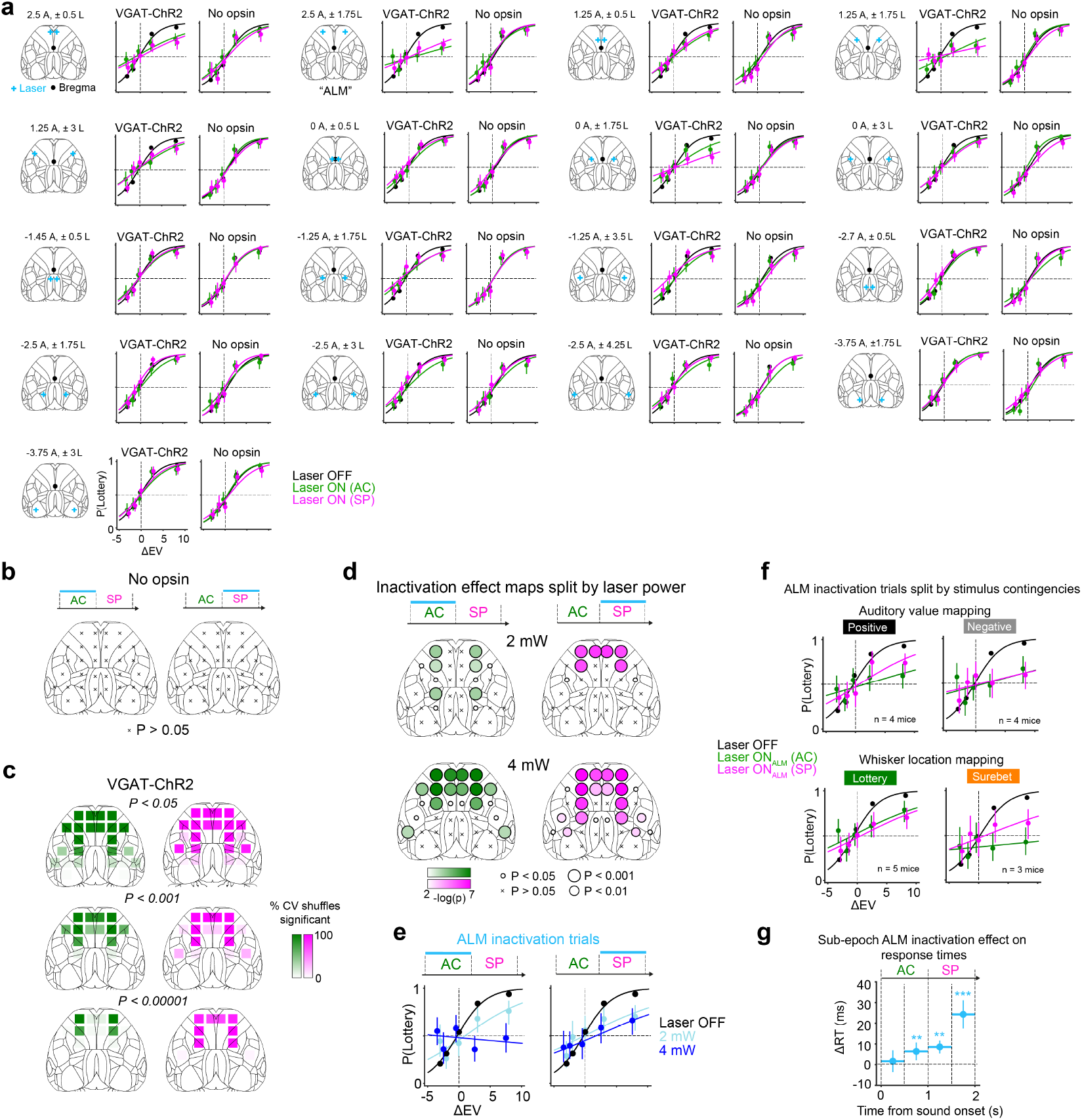
Optogenetic effects on task performance across regions, laser powers and stimulus-contingency groups. **a**, Psychometric curves for all dorsal cortical sites targeted for bilateral photoinhibition. For each region: Left, photoinhibition target locations (blue) and stereotaxic coordinates relative to bregma (black). Middle, behavioural performance of VGAT-ChR2 mice (n = 8), showing control trials (black) and trials with photoinhibition during either the abstract choice epoch (AC; 0 − 1 s; green) or spatial planning epoch (SP; 1 − 2 s; magenta). Right, corresponding data for ‘no opsin’ control mice (n = 6). **b**, Statistical significance of region and epoch-specific optogenetics experiments in control mice, quantified using cross-validated (CV) generalised linear mixed-effects models (GLMMs). Dorsal cortex maps show the median *P* value across 50 CV shuffles corrected for 17 regional comparisons. ‘x’ markers indicate *P* > 0.05. **c**, Reliability of statistical effects across CV shuffles. Colour scale indicates the percentage of iterations that met the *P* value threshold indicated. Left, VGAT-ChR2 mice (n = 8). **d**, Similar to **b**, but showing behavioural effects in VGAT-ChR2 mice separated by 2 vs 4 mW laser power trials. **e**, Psychometric curves from 2 vs 4 mW ALM photoinhibition trials. **f**, ALM photoinhibition effects across different mice grouped by stimulus contingency. Top row, psychometric curves split by auditory-to-value mapping. Bottom row, psychometric curves split by whisker-to-offer location mapping. Colour code same as **a. g**, Effect of sub-epoch ALM photoinhibition on response times (RTs). The horizontal blue bars indicate the timing and duration of each photoinhibition sub-epoch trial-type, circular markers show median RTs, vertical errorbars show s.e.m across trials. Data from 5 VGAT-ChR2-EYFP mice. Paired comparisons to control trials used Wilcoxon ranksum tests. Significance is indicated as \**P* < 0.05, \*\**P* < 0.01, \*\*\**P* < 0.001; ns, non-significant (*P* > 0.05).

**Fig. S8:**
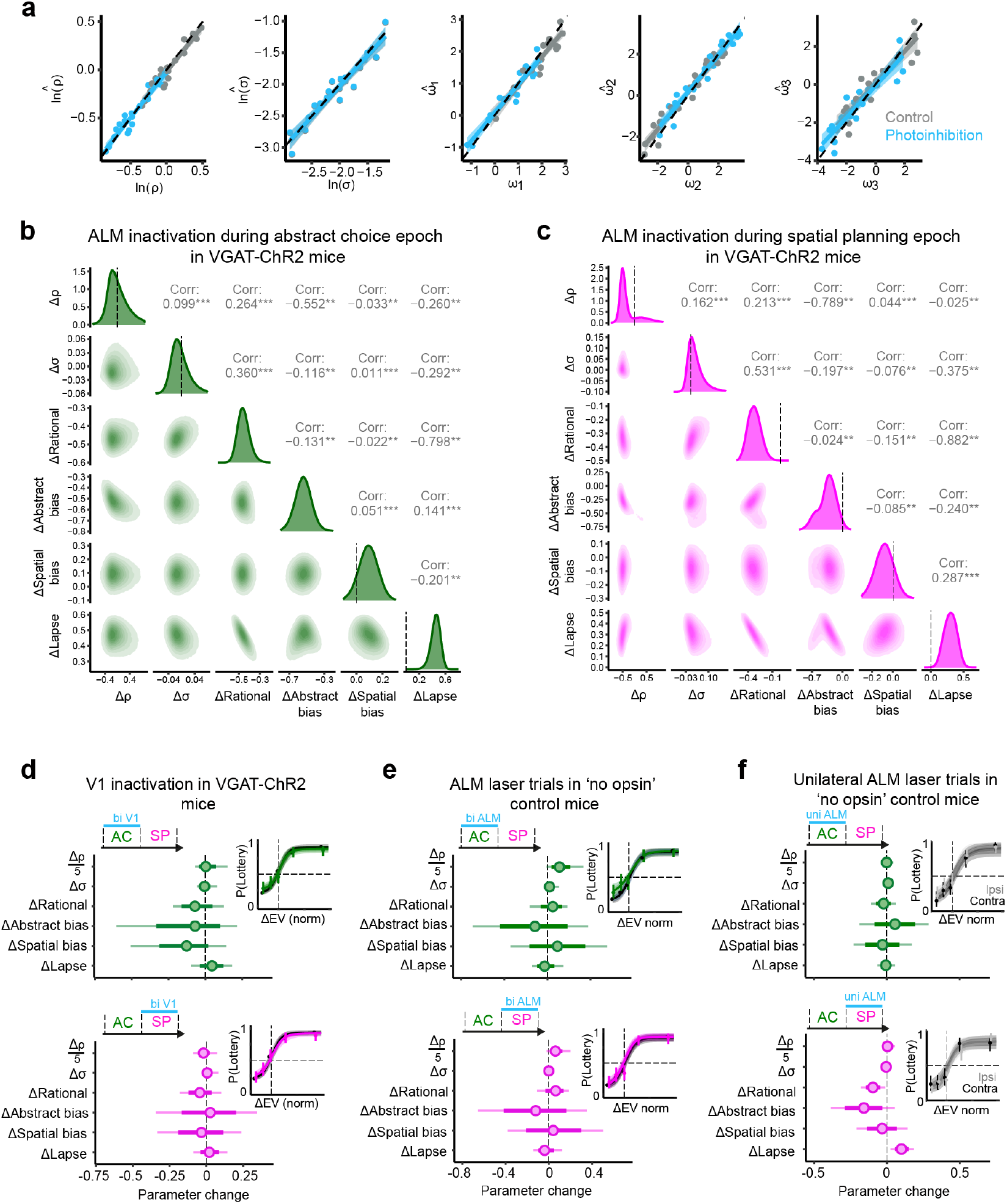
Mixed-agent model validation and fits to control data. **a**, Mixed-agent model validation using synthetic data. True and recovered parameters are plotted. Individual points show simulated data from 20 subjects. Control and photoinhibition conditions are shown in grey and blue respectively. Lines indicate linear model fits across subjects. Simulated optogenetic effects: Δ ln *ρ* = − 0.5, Δ*ω*_1_ = − 1, Δ*ω*_2_ = +1, Δ*ω*_3_ = − 1. **b**, 2D marginal posteriors show minimal trade-offs between pairs of model parameters, confirming well-identified estimates for each parameter for bilateral ALM inhibition during the abstract choice epoch (n = 8 VGAT-ChR2 mice). Correlation coefficient (Pearson’s r) and significance is indicated as \**P* < 0.05, \*\**P* < 0.01, \*\*\**P* < 0.001. **c**, Similar to **b**, for ALM inhibition during the spatial planning epoch. **d**, Mixed-agent model fits to bilateral V1 photoinhibition in VGAT-ChR2 mice (n = 8), for first epoch (top) and second epoch (bottom) photoinhibition. Insets, psychometric model fits to the choice data. **e**, Similar to **d**, but for bilateral ALM laser delivery trials in control mice without ChR2 expression (n = 6). **f**, Similar to **d**, but for unilateral ALM laser delivery trials in control mice without ChR2 expression (n = 6).

**Fig. S9:**
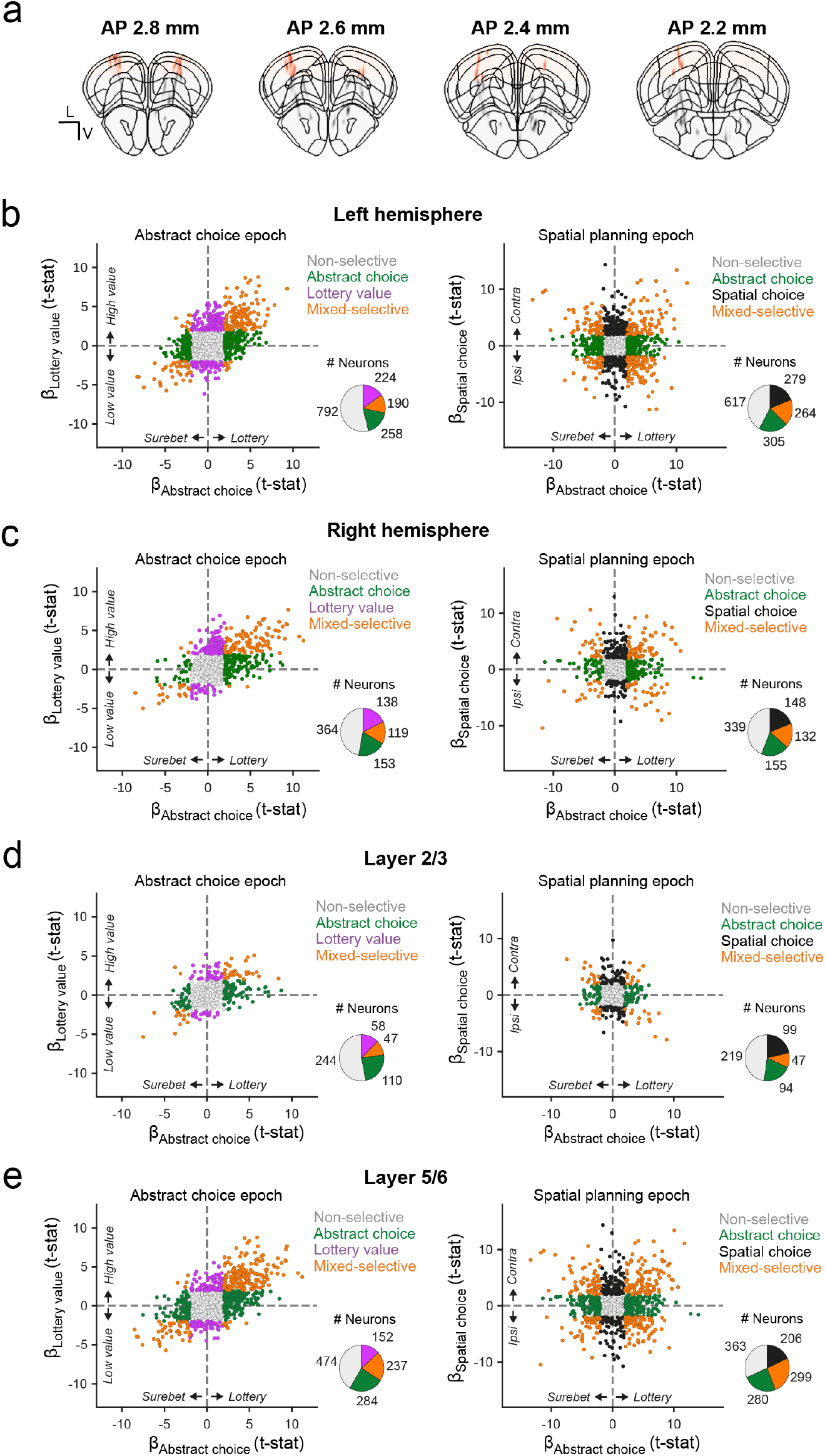
Value-to-action coding in ALM across recording hemispheres and cortical layers. **a**, Neuropixels probe tract locations for 4 recording mice, coloured tracks highlight parts of the probes within ALM. **b**, Task selectivity for all ALM neurons recorded in the left hemisphere. Left, coefficients of lottery value versus abstract choice regressors during the first epoch (0 − 1 s). Right, coefficients of spatial choice (contra- or ipsilateral to the recording site) vs abstract choice regressors during the second epoch (1 − 2 s). Each point represents an individual neuron; colours indicate significantly selective neurons. Inset pie chart summarizes the number and proportion of selective neurons. **c**, Similar to **b**, but showing results from all ALM neurons recorded in the right hemisphere. **d**, Similar to **b**, but showing results from ALM neurons in layers 2 and 3 (superficial layers). **e**, Similar to **d**, but showing results from ALM neurons in layers 5 and 6 (deep layers).

**Fig. S10:**
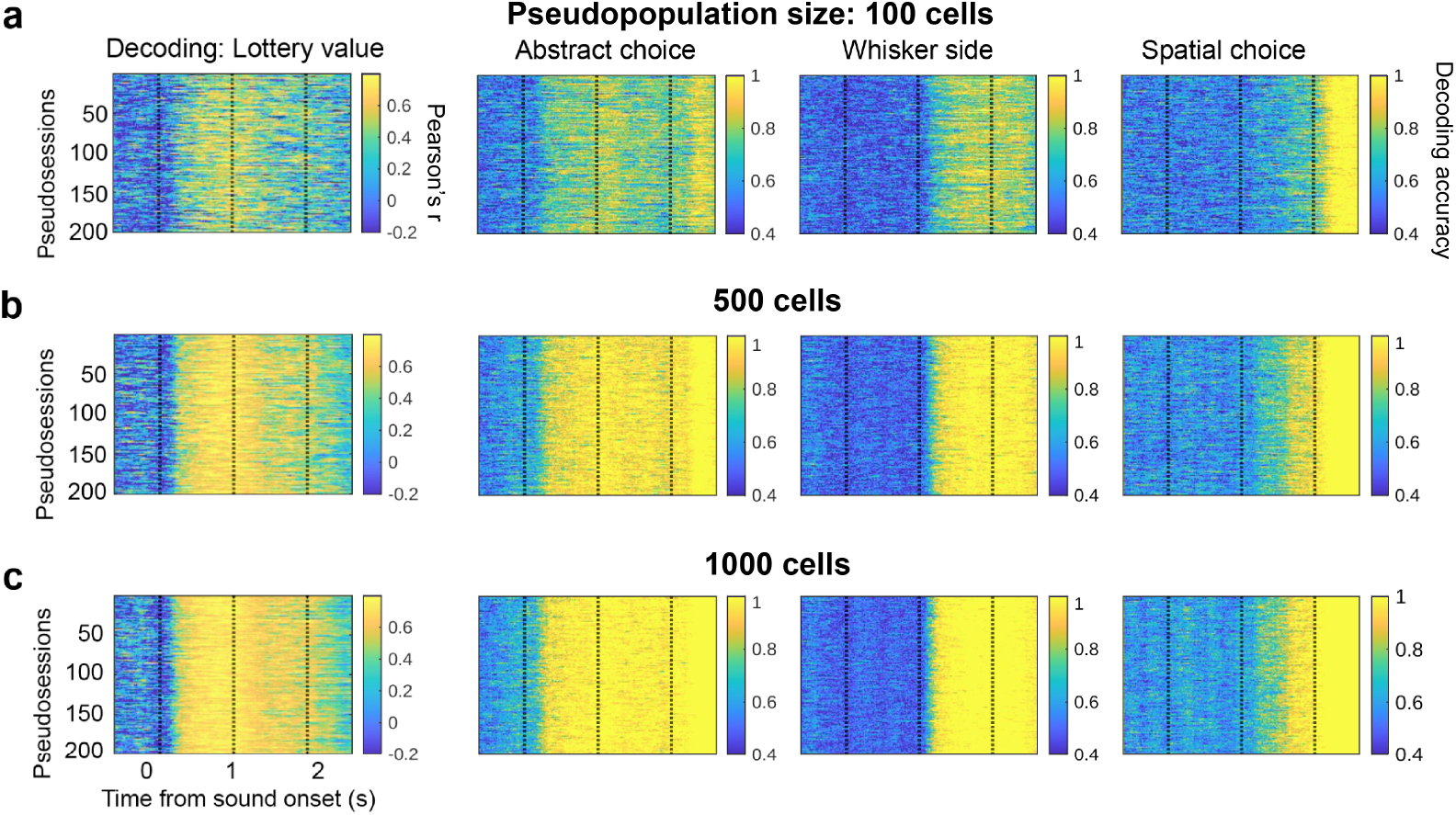
Population decoding for all pseudosessions. **a**, Similar to Fig. 4e-h, for all 200 pseudosessions. Population decoding performance of lottery value, abstract (lottery versus surebet) choices, whisker stimulus sides, and spatial (left versus right) choices over time, for pseudopopulation size of 100 cells. *P* values were calculated for each pseudosession, and median *P* was reported in Fig. 4e-h. **b**, Similar to **a**, for pseudopopulation size of 500 cells. **c**, Similar to **a**, for pseudopopulation size of 1000 cells.

## References

[1] Matthew P.H. Gardner, Davied Sanchez, Jessica C. Conroy, Andrew M. Wikenheiser, Jingfeng Zhou, and Geoffrey Schoenbaum. Processing in Lateral Orbitofrontal Cortex Is Required to Estimate Subjective Preference during Initial, but Not Established, Economic Choice. Neuron, 108(3):526–537.e4, November 2020. doi: 10.1016/j.neuron.2020.08.010.

[2] Christine M. Constantinople, Alex T. Piet, and Carlos D. Brody. An Analysis of Decision under Risk in Rats. Current Biology, 29(12):2066–2074.e5, June 2019. doi: 10.1016/j.cub.2019.05.013.

[3] Alireza Soltani and Alicia Izquierdo. Adaptive learning under expected and unexpected uncertainty. Nature Reviews Neuroscience, 20(10):635–644, October 2019. doi: 10.1038/s41583-019-0180-y.

[4] Xinying Cai, Soyoun Kim, and Daeyeol Lee. Heterogeneous Coding of Temporally Discounted Values in the Dorsal and Ventral Striatum during Intertemporal Choice. Neuron, 69(1):170– 182, January 2011. doi: 10.1016/j.neuron.2010.11.041.

[5] Evgeniya Lukinova, Yuyue Wang, Steven F Lehrer, and Jeffrey C Erlich. Time preferences are reliable across time-horizons and verbal versus experiential tasks. eLife, 8:e39656, February 2019. doi: 10.7554/eLife.39656.

[6] Michael Inzlicht, Amitai Shenhav, and Christopher Y. Olivola. The Effort Paradox: Effort Is Both Costly and Valued. Trends in Cognitive Sciences, 22(4):337–349, April 2018. doi: 10.1016/j.tics.2018.01.007.

[7] Xinying Cai and Camillo Padoa-Schioppa. Neuronal activity in dorsal anterior cingulate cortex during economic choices under variable action costs. eLife, 10:e71695, October 2021. doi: 10.7554/eLife.71695.

[8] 23and Me Research Team, eQTLgen Consortium, International Cannabis Consortium, Social Science Genetic Association Consortium, Richard Karlsson Linnér, Pietro Biroli, Edward Kong, S. Fleur W. Meddens, Robbee Wedow, Mark Alan Fontana, Maël Lebreton, Stephen P. Tino, Abdel Abdellaoui, Anke R. Hammerschlag, Michel G. Nivard, Aysu Okbay, Cornelius A. Rietveld, Pascal N. Timshel, Maciej Trzaskowski, Ronald de Vlaming, Christian L. Zünd, Yanchun Bao, Laura Buzdugan, Ann H. Caplin, Chia-Yen Chen, Peter Eibich, Pierre Fontanillas, Juan R. Gonzalez, Peter K. Joshi, Ville Karhunen, Aaron Kleinman, Remy Z. Levin, Christina M. Lill, Gerardus A. Meddens, Gerard Muntané, Sandra Sanchez-Roige, Frank J. van Rooij, Erdogan Taskesen, Yang Wu, Futao Zhang, Adam Auton, Jason D. Boardman, David W. Clark, Andrew Conlin, Conor C. Dolan, Urs Fischbacher, Patrick J. F. Groenen, Kathleen Mullan Harris, Gregor Hasler, Albert Hofman, Mohammad A. Ikram, Sonia Jain, Robert Karlsson, Ronald C. Kessler, Maarten Kooyman, James MacKillop, Minna Männikkö, Carlos Morcillo-Suarez, Matthew B. McQueen, Klaus M. Schmidt, Melissa C. Smart, Matthias Sutter, A. Roy Thurik, André G. Uitterlinden, Jon White, Harriet de Wit, Jian Yang, Lars Bertram, Dorret I. Boomsma, Tõnu Esko, Ernst Fehr, David A. Hinds, Magnus Johannesson, Meena Kumari, David Laibson, Patrik K. E. Magnusson, Michelle N. Meyer, Arcadi Navarro, Abraham A. Palmer, Tune H. Pers, Danielle Posthuma, Daniel Schunk, Murray B. Stein, Rauli Svento, Henning Tiemeier, Paul R. H. J. Timmers, Patrick Turley, Robert J. Ursano, Gert G. Wagner, James F. Wilson, Jacob Gratten, James J. Lee, David Cesarini, Daniel J. Benjamin, Philipp D. Koellinger, and Jonathan P. Beauchamp. Genome-wide association analyses of risk tolerance and risky behaviors in over 1 million individuals identify hundreds of loci and shared genetic influences. Nature Genetics, 51(2):245–257, February 2019. doi: 10.1038/s41588-018-0309-3.

[9] the 23andMe Research Team, Sandra Sanchez-Roige, Pierre Fontanillas, Sarah L. Elson, Anita Pandit, Ellen M. Schmidt, Johanna R. Foerster, Gonçalo R. Abecasis, Joshua C. Gray, Harriet de Wit, Lea K. Davis, James MacKillop, and Abraham A. Palmer. Genome-wide association study of delay discounting in 23,217 adult research participants of European ancestry. Nature Neuroscience, December 2017. doi: 10.1038/s41593-017-0032-x.

[10] Camillo Padoa-Schioppa. Neurobiology of Economic Choice: A Good-Based Model. Annual Review of Neuroscience, 34(1):333–359, July 2011. doi: 10.1146/annurev-neuro-061010-113648.

[11] Paul Cisek and John F. Kalaska. Neural Mechanisms for Interacting with a World Full of Action Choices. Annual Review of Neuroscience, 33(1):269–298, June 2010. doi: 10.1146/annurev.neuro.051508.135409.

[12] Antonio Rangel and Todd Hare. Neural computations associated with goal-directed choice. Current Opinion in Neurobiology, 20(2):262–270, April 2010. doi: 10.1016/j.conb.2010.03.001.

[13] Camillo Padoa-Schioppa and John A. Assad. Neurons in the orbitofrontal cortex encode economic value. Nature, 441(7090):223–226, May 2006. doi: 10.1038/nature04676.

[14] Beizhen Zhang, Janis Ying Ying Kan, Mingpo Yang, Xiaochun Wang, Jiahao Tu, and Michael Christopher Dorris. Transforming absolute value to categorical choice in primate superior colliculus during value-based decision making. Nature Communications, 12(1):3410, June 2021. doi: 10.1038/s41467-021-23747-z.

[15] L. Ding and O. Hikosaka. Comparison of Reward Modulation in the Frontal Eye Field and Caudate of the Macaque. Journal of Neuroscience, 26(25):6695–6703, June 2006. doi: 10.1523/JNEUROSCI.0836-06.2006.

[16] Michael L. Platt and Paul W. Glimcher. Neural correlates of decision variables in parietal cortex. Nature, 400(6741):233–238, July 1999. doi: 10.1038/22268.

[17] Soyoun Kim, Jaewon Hwang, and Daeyeol Lee. Prefrontal Coding of Temporally Discounted Values during Intertemporal Choice. Neuron, 59(1):161–172, July 2008. doi: 10.1016/j.neuron.2008.05.010.

[18] Brian Lau and Paul W. Glimcher. Value Representations in the Primate Striatum during Matching Behavior. Neuron, 58(3):451–463, May 2008. doi: 10.1016/j.neuron.2008.02.021.

[19] Paul Cisek. Making decisions through a distributed consensus. Current Opinion in Neurobiology, 22(6):927–936, December 2012. doi: 10.1016/j.conb.2012.05.007.

[20] Xinying Cai and Camillo Padoa-Schioppa. Contributions of Orbitofrontal and Lateral Prefrontal Cortices to Economic Choice and the Good-to-Action Transformation. Neuron, 81(5):1140–1151, March 2014. doi: 10.1016/j.neuron.2014.01.008.

[21] Chaofei Bao, Xiaoyue Zhu, Joshua Mōller-Mara, Jingjie Li, Sylvain Dubroqua, and Jeffrey C. Erlich. The rat frontal orienting field dynamically encodes value for economic decisions under risk. Nature Neuroscience, 26(11):1942–1952, November 2023. doi: 10.1038/s41593-023-01461-x.

[22] Masaru Kuwabara, Ningdong Kang, Timothy E Holy, and Camillo Padoa-Schioppa. Neural mechanisms of economic choices in mice. eLife, 9:e49669, February 2020. doi: 10.7554/eLife.49669.

[23] Matthew P. H. Gardner, Jessica S. Conroy, Michael H. Shaham, Clay V. Styer, and Geoffrey Schoenbaum. Lateral Orbitofrontal Inactivation Dissociates Devaluation-Sensitive Behavior and Economic Choice. Neuron, 96(5):1192–1203.e4, December 2017. doi: 10.1016/j.neuron.2017.10.026.

[24] Christine M Constantinople, Alex T Piet, Peter Bibawi, Athena Akrami, Charles Kopec, and Carlos D Brody. Lateral orbitofrontal cortex promotes trial-by-trial learning of risky, but not spatial, biases. eLife, 8:e49744, November 2019. doi: 10.7554/eLife.49744.

[25] Antara Majumdar, Caitlin Ashcroft, Matthias Fritsche, Sandra Tan, Peter Zatka-Haas, Orsolya Folsz, Niamh Walker, Leah Mistry, Anita M. Rominto, Marko Tvrdic, Zoltán Molnár, Huriye Atilgan, Adam M. Packer, Simon J.B. Butt, and Armin Lak. Distinct representations of economic variables across regions and projections of the frontal cortex. Neuron, page S0896627325007159, October 2025. doi: 10.1016/j.neuron.2025.09.027.

[26] Alexandra Stolyarova and Alicia Izquierdo. Complementary contributions of basolateral amygdala and orbitofrontal cortex to value learning under uncertainty. eLife, 6:e27483, July 2017. doi: 10.7554/eLife.27483.

[27] Nicole L. Jenni, Debra A. Bercovici, and Stan B. Floresco. Medial Orbitofrontal, Prefrontal, and Amygdalar Circuits Support Dissociable Component Processes of Risk/Reward Decision-Making. The Journal of Neuroscience, 45(16):e2147242025, April 2025. doi: 10.1523/JNEUROSCI.2147-24.2025.

[28] Jackson D. Schumacher, Mieke Van Holstein, Peiran Zhou, Paula E. MacLeod, Vaishali Bagrodia, and Stan B. Floresco. Complementary yet dissociable influences of medial and lateral orbitofrontal cortex over cue-guided decisions involving reward magnitude and uncertainty. The Journal of Neuroscience, page e1989242025, October 2025. doi: 10.1523/JNEUROSCI.1989-24.2025.

[29] Hidehiko K. Inagaki, Susu Chen, Kayvon Daie, Arseny Finkelstein, Lorenzo Fontolan, Sandro Romani, and Karel Svoboda. Neural Algorithms and Circuits for Motor Planning. Annual Review of Neuroscience, 45(1):249–271, July 2022. doi: 10.1146/annurev-neuro-092021-121730.

[30] Zengcai V. Guo, Nuo Li, Daniel Huber, Eran Ophir, Diego Gutnisky, Jonathan T. Ting, Guoping Feng, and Karel Svoboda. Flow of Cortical Activity Underlying a Tactile Decision in Mice. Neuron, 81(1):179–194, January 2014. doi: 10.1016/j.neuron.2013.10.020.

[31] Nuo Li, Tsai-Wen Chen, Zengcai V. Guo, Charles R. Gerfen, and Karel Svoboda. A motor cortex circuit for motor planning and movement. Nature, 519(7541):51–56, March 2015. doi: 10.1038/nature14178.

[32] Chunyu A. Duan, Yuxin Pan, Guofen Ma, Taotao Zhou, Siyu Zhang, and Ning-long Xu. A cortico-collicular pathway for motor planning in a memory-dependent perceptual decision task. Nature Communications, 12(1):2727, May 2021. doi: 10.1038/s41467-021-22547-9.

[33] Zoe C. Ashwood, Nicholas A. Roy, Iris R. Stone, The International Brain Laboratory, Anne E. Urai, Anne K. Churchland, Alexandre Pouget, and Jonathan W. Pillow. Mice alternate between discrete strategies during perceptual decision-making. Nature Neuroscience, 25(2): 201–212, February 2022. doi: 10.1038/s41593-021-01007-z.

[34] Scott S. Bolkan, Iris R. Stone, Lucas Pinto, Zoe C. Ashwood, Jorge M. Iravedra Garcia, Alison L. Herman, Priyanka Singh, Akhil Bandi, Julia Cox, Christopher A. Zimmerman, Jounhong Ryan Cho, Ben Engelhard, Jonathan W. Pillow, and Ilana B. Witten. Opponent control of behavior by dorsomedial striatal pathways depends on task demands and internal state. Nature Neuroscience, 25(3):345–357, March 2022. doi: 10.1038/s41593-022-01021-9.

[35] Simon Musall, Matthew T. Kaufman, Ashley L. Juavinett, Steven Gluf, and Anne K. Churchland. Single-trial neural dynamics are dominated by richly varied movements. Nature Neuroscience, 22(10):1677–1686, October 2019. doi: 10.1038/s41593-019-0502-4.

[36] Michael Lohse, Oliver M Gauld, Maja T Skrętowska, Peter Vincent, Quentin Pajot-Moric, Simon Townsend, Chunyu A Duan, Athena Akrami, Thomas D Mrsic-Flogel, and Robert AA Campbell. Zapit: Open Source Random-Access Photostimulation For Neuroscience. bioRxiv, February 2024. doi: 10.1101/2024.02.12.579892.

[37] Lucas Pinto, Kanaka Rajan, Brian DePasquale, Stephan Y. Thiberge, David W. Tank, and Carlos D. Brody. Task-Dependent Changes in the Large-Scale Dynamics and Necessity of Cortical Regions. Neuron, 104(4):810–824.e9, November 2019. doi: 10.1016/j.neuron.2019.08.025.

[38] Nuo Li, Susu Chen, Zengcai V Guo, Han Chen, Yan Huo, Hidehiko K Inagaki, Guang Chen, Courtney Davis, David Hansel, Caiying Guo, and Karel Svoboda. Spatiotemporal constraints on optogenetic inactivation in cortical circuits. eLife, 8:e48622, November 2019. doi: 10.7554/eLife.48622.

[39] Jia Shen, Nuttida Rungratsameetaweemana, Prayshita Sharma, Darcy S. Peterka, Herbert Zheng Wu, and Michael N. Shadlen. ALM enables contextual decision-making via dynamic reconfiguration of local circuits. bioRxiv, September 2025. doi: 10.1101/2025.09.11.675319.

[40] Aldo Rustichini and Camillo Padoa-Schioppa. A neuro-computational model of economic decisions. Journal of Neurophysiology, 114(3):1382–1398, September 2015. doi: 10.1152/jn.00184.2015.

[41] Aldo Battista, Camillo Padoa-Schioppa, and Xiao-Jing Wang. A neural circuit framework for economic choice: From building blocks of valuation to compositionality in multitasking. bioRxiv, March 2025. doi: 10.1101/2025.03.13.643098.

[42] Man Yi Yim, Xinying Cai, and Xiao-Jing Wang. Transforming the Choice Outcome to an Action Plan in Monkey Lateral Prefrontal Cortex: A Neural Circuit Model. Neuron, 103(3):520–532.e5, August 2019. doi: 10.1016/j.neuron.2019.05.032.

[43] Chunyu A Duan, Jeffrey C Erlich, and Carlos D Brody. Requirement of Prefrontal and Midbrain Regions for Rapid Executive Control of Behavior in the Rat. Neuron, 86(6):1491– 1503, June 2015. doi: 10.1016/j.neuron.2015.05.042.

[44] Sharath Bennur and Joshua I. Gold. Distinct Representations of a Perceptual Decision and the Associated Oculomotor Plan in the Monkey Lateral Intraparietal Area. The Journal of Neuroscience, 31(3):913–921, January 2011. doi: 10.1523/JNEUROSCI.4417-10.2011.

[45] Shinichiro Kira, Houman Safaai, Ari S. Morcos, Stefano Panzeri, and Christopher D. Harvey. A distributed and efficient population code of mixed selectivity neurons for flexible navigation decisions. Nature Communications, 14(1):2121, April 2023. doi: 10.1038/s41467-023-37804-2.

[46] Gouki Okazawa and Roozbeh Kiani. Neural Mechanisms That Make Perceptual Decisions Flexible. Annual Review of Physiology, 85(1):191–215, February 2023. doi: 10.1146/annurev-physiol-031722-024731.

[47] S. Shushruth, Ariel Zylberberg, and Michael N. Shadlen. Sequential sampling from memory underlies action selection during abstract decision-making. Current Biology, 32(9):1949– 1960.e5, May 2022. doi: 10.1016/j.cub.2022.03.014.

[48] Chunyu A. Duan, Marino Pagan, Alex T. Piet, Charles D. Kopec, Athena Akrami, Alexander J. Riordan, Jeffrey C. Erlich, and Carlos D. Brody. Collicular circuits for flexible sensorimotor routing. Nature Neuroscience, 24(8):1110–1120, August 2021. doi: 10.1038/s41593-021-00865-x.

[49] Ivana Orsolic, Maxime Rio, Thomas D. Mrsic-Flogel, and Petr Znamenskiy. Mesoscale cortical dynamics reflect the interaction of sensory evidence and temporal expectation during perceptual decision-making. Neuron, 109(11):1861–1875.e10, June 2021. doi: 10.1016/j.neuron.2021.03.031.

[50] Daniel Hulsey, Kevin Zumwalt, Luca Mazzucato, David A. McCormick, and Santiago Jaramillo. Decision-making dynamics are predicted by arousal and uninstructed movements. Cell Reports, 43(2):113709, February 2024. doi: 10.1016/j.celrep.2024.113709.

[51] Carsen Stringer, Marius Pachitariu, Nicholas Steinmetz, Charu Bai Reddy, Matteo Carandini, and Kenneth D. Harris. Spontaneous behaviors drive multidimensional, brainwide activity. Science, 364(6437):eaav7893, April 2019. doi: 10.1126/science.aav7893.

[52] Philip Coen, Timothy P.H. Sit, Miles J. Wells, Matteo Carandini, and Kenneth D. Harris. Mouse frontal cortex mediates additive multisensory decisions. Neuron, 111(15):2432–2447.e13, August 2023. doi: 10.1016/j.neuron.2023.05.008.

[53] Zengcai V. Guo, Hidehiko K. Inagaki, Kayvon Daie, Shaul Druckmann, Charles R. Gerfen, and Karel Svoboda. Maintenance of persistent activity in a frontal thalamocortical loop. Nature, 545(7653):181–186, May 2017. doi: 10.1038/nature22324.

[54] Susu Chen, Yi Liu, Ziyue Aiden Wang, Jennifer Colonell, Liu D. Liu, Han Hou, Nai-Wen Tien, Tim Wang, Timothy Harris, Shaul Druckmann, Nuo Li, and Karel Svoboda. Brain-wide neural activity underlying memory-guided movement. Cell, 187(3):676–691.e16, February 2024. doi: 10.1016/j.cell.2023.12.035.

[55] Yu Wang, Xinxin Yin, Zhouzhou Zhang, Jiejue Li, Wenyu Zhao, and Zengcai V. Guo. A cortico-basal ganglia-thalamo-cortical channel underlying short-term memory. Neuron, 109 (21):3486–3499.e7, November 2021. doi: 10.1016/j.neuron.2021.08.002.

[56] Nuo Li, Kayvon Daie, Karel Svoboda, and Shaul Druckmann. Robust neuronal dynamics in premotor cortex during motor planning. Nature, 532(7600):459–464, April 2016. doi: 10.1038/nature17643.

[57] Xinxin Yin, Yu Wang, Jiejue Li, and Zengcai V. Guo. Lateralization of short-term memory in the frontal cortex. Cell Reports, 40(7):111190, August 2022. doi: 10.1016/j.celrep.2022.111190.

[58] Daniel J. O’Shea, Lea Duncker, Werapong Goo, Xulu Sun, Saurabh Vyas, Eric M. Trautmann, Ilka Diester, Charu Ramakrishnan, Karl Deisseroth, Maneesh Sahani, and Krishna V. Shenoy. Direct neural perturbations reveal a dynamical mechanism for robust computation. bioRxiv, December 2022. doi: 10.1101/2022.12.16.520768.

[59] Michael N. Economo, Sarada Viswanathan, Bosiljka Tasic, Erhan Bas, Johan Winnubst, Vilas Menon, Lucas T. Graybuck, Thuc Nghi Nguyen, Kimberly A. Smith, Zizhen Yao, Lihua Wang, Charles R. Gerfen, Jayaram Chandrashekar, Hongkui Zeng, Loren L. Looger, and Karel Svoboda. Distinct descending motor cortex pathways and their roles in movement. Nature, 563(7729):79–84, November 2018. doi: 10.1038/s41586-018-0642-9.

[60] Lea Duncker and Maneesh Sahani. Dynamics on the manifold: Identifying computational dynamical activity from neural population recordings. Current Opinion in Neurobiology, 70: 163–170, October 2021. doi: 10.1016/j.conb.2021.10.014.

[61] Jae-Hyun Kim, Kayvon Daie, and Nuo Li. A combinatorial neural code for long-term motor memory. Nature, 637(8046):663–672, January 2025. doi: 10.1038/s41586-024-08193-3.

[62] Xia Chen, Xiao Yao, Xinxin Yin, and Zengcai V. Guo. Task-similarity dependent reconfiguration of compositional modules and geometry in frontal cortex. bioRxiv, November 2025. doi: 10.1101/2025.11.06.687080.

[63] Zheng Wu, Ashok Litwin-Kumar, Philip Shamash, Alexei Taylor, Richard Axel, and Michael N. Shadlen. Context-Dependent Decision Making in a Premotor Circuit. Neuron, 106(2):316–328.e6, April 2020. doi: 10.1016/j.neuron.2020.01.034.

[64] Arash Bellafard, Ghazal Namvar, Jonathan C. Kao, Alipasha Vaziri, and Peyman Golshani. Volatile working memory representations crystallize with practice. Nature, 629(8014):1109– 1117, May 2024. doi: 10.1038/s41586-024-07425-w.

[65] Talmo D. Pereira, Nathaniel Tabris, Arie Matsliah, David M. Turner, Junyu Li, Shruthi Ravindranath, Eleni S. Papadoyannis, Edna Normand, David S. Deutsch, Z. Yan Wang, Grace C. McKenzie-Smith, Catalin C. Mitelut, Marielisa Diez Castro, John D’Uva, Mikhail Kislin, Dan H. Sanes, Sarah D. Kocher, Samuel S.-H. Wang, Annegret L. Falkner, Joshua W. Shaevitz, and Mala Murthy. SLEAP: A deep learning system for multi-animal pose tracking. Nature Methods, 19(4):486–495, April 2022. doi: 10.1038/s41592-022-01426-1.

[66] Joao Couto, Simon Musall, Xiaonan R. Sun, Anup Khanal, Steven Gluf, Shreya Saxena, Ian Kinsella, Taiga Abe, John P. Cunningham, Liam Paninski, and Anne K. Churchland. Chronic, cortex-wide imaging of specific cell populations during behavior. Nature Protocols, 16(7):3241–3263, July 2021. doi: 10.1038/s41596-021-00527-z.

[67] G. N. Wilkinson and C. E. Rogers. Symbolic Description of Factorial Models for Analysis of Variance. Applied Statistics, 22(3):392, 1973. doi: 10.2307/2346786.

[68] Philippe N. Tobler and Elke U. Weber. Valuation for Risky and Uncertain Choices. Neuroeconomics: Decision Making and the Brain: Second Edition, pages 149–172, September 2013. doi: 10.1016/B978-0-12-416008-8.00009-7.

[69] Paul-Christian Bürkner. Brms: An R Package for Bayesian Multilevel Models Using Stan. Journal of Statistical Software, 80(1), 2017. doi: 10.18637/jss.v080.i01.

[70] Stan Development Team. RStan: The R interface to Stan, 2025.

[71] Marius Pachitariu, Shashwat Sridhar, Jacob Pennington, and Carsen Stringer. Spike sorting with Kilosort4. Nature Methods, 21(5):914–921, May 2024. doi: 10.1038/s41592-024-02232-7.

[72] Alessio P Buccino, Cole L Hurwitz, Samuel Garcia, Jeremy Magland, Joshua H Siegle, Roger Hurwitz, and Matthias H Hennig. SpikeInterface, a unified framework for spike sorting. eLife, 9:e61834, November 2020. doi: 10.7554/eLife.61834.

[73] Andrei Khilkevich, Michael Lohse, Ryan Low, Ivana Orsolic, Tadej Bozic, Paige Windmill, and Thomas D. Mrsic-Flogel. Brain-wide dynamics linking sensation to action during decisionmaking. Nature, 634(8035):890–900, October 2024. doi: 10.1038/s41586-024-07908-w.

[74] Charles R. Harris, K. Jarrod Millman, Stéfan J. van der Walt, Ralf Gommers, Pauli Virtanen, David Cournapeau, Eric Wieser, Julian Taylor, Sebastian Berg, Nathaniel J. Smith, Robert Kern, Matti Picus, Stephan Hoyer, Marten H. van Kerkwijk, Matthew Brett, Allan Haldane, Jaime Fernández del Río, Mark Wiebe, Pearu Peterson, Pierre Gérard-Marchant, Kevin Sheppard, Tyler Reddy, Warren Weckesser, Hameer Abbasi, Christoph Gohlke, and Travis E. Oliphant. Array programming with NumPy. Nature, 585(7825):357–362, September 2020. doi: 10.1038/s41586-020-2649-2.

